# Dosage-sensitive *RBFOX2* autoregulation promotes cardiomyocyte differentiation by maturing the transcriptome

**DOI:** 10.1101/2025.10.28.685214

**Authors:** Mengmeng Huang, Feria A. Ladha, Yunxia Wang, Michael A. Trembley, Hui-Min Yin, Rongbin Zheng, Maksymilian Prondzynski, Yashavi Tharani, Alexander A. Akerberg, Stefan Aigner, Brian A. Yee, Joshua Mayourian, Sarah U. Morton, Vassilios J. Bezzerides, William T. Pu, Gene W. Yeo, Kaifu Chen, C. Geoffrey Burns, Caroline E. Burns

## Abstract

Haploinsufficiency of the RNA splicing regulator, *RBFOX2*, is linked to congenital heart disease (CHD), yet its pathogenic mechanisms remain unclear. Here, we demonstrate that RBFOX2 is essential for progressing cardiomyocyte (CM) differentiation by shifting exon usage profiles to more mature patterns in sarcomere, cytoskeletal, and focal adhesion genes, including *alpha-actinin-2* (*ACTN2*). This maturation program is initiated by critical levels of RBFOX2 that facilitate autoregulatory splicing at mutually exclusive exons encoding early and late isoforms with distinct functional roles. In heterozygous CMs, autoregulation is disrupted, which skews isoform ratios and generates a dominant-negative product caused by exon co-inclusion. Finally, we demonstrate that overexpression of ACTN2 rescues heterozygous, but not null, phenotypes by restoring contractility, which triggers a mechanosensing feedback loop involving upregulation of RBFOX2 from the wildtype allele and transcriptome maturation. Our data suggest that decreased RBFOX2 dosage and autoregulation impair CM differentiation, contributing to CHD pathogenesis and heart failure susceptibility.

## INTRODUCTION

Congenital heart disease (CHD) is among the most common birth defects, affecting ∼1% of live births and accounting for nearly 40% of childhood morbidity from congenital anomalies^1,2^. Although advances in prenatal diagnosis, surgical palliation, and perioperative management have improved survival, ventricular dysfunction that progresses to heart failure (HF) remains a major cause of mortality across the life span^3,4^.

This risk is particularly high in complex and heterogenous CHD subtypes such as hypoplastic left heart syndrome (HLHS) where left-sided structures, including the left ventricle, aorta, and valves, are variably underdeveloped^5^. Standard care involves three staged palliative surgeries to reroute the circulation for a single ventricle (SV) physiology, with the first two occurring in the first year of life. In this first year, when the balance between the systemic and pulmonary circulations is unstable, nearly 34% of infants die or require a heart transplant^6–8^. The third surgical stage, the Fontan procedure, occurs at 2-4 years of age^9^. In these patients, the need for transplant remains high, primarily due ventricular dysfunction that often progresses to HF^8,10^. In fact, only about half of people with HLHS remain transplant-free by 10-years of age, with the other half experiencing death or transplant^8,11–13^. These divergent outcomes suggest that additional factors, including genetics, influence HF susceptibility in the setting of altered hemodynamics. Moreover, patients surviving with other forms of CHD also show this differential pattern of HF susceptibility^14^. As such, new mechanistic insights into the genetic causes of CHD, and their impact on ventricular dysfunction and HF outcomes, are needed for improved risk stratification and targeted clinical intervention.

One gene associated with CHD, including HLHS, is *RBFOX2*^15,16^, which encodes a highly conserved RNA binding protein with known roles in alterative splicing^17^. In a cohort of 11,555 probands, 74 individuals carried heterozygous *RBFOX2* variants with predicted damaging functions across diverse CHD subtypes, including 10 with HLHS^16^. Although the overall number was too low to reach genome-wide statistical significance, participants with HLHS were disproportionately enriched for protein-truncating or protein-damaging variants, suggesting a link between HLHS and *RBFOX2* haploinsufficiency.

As observed with many CHD genes, this dosage sensitivity appears unique to humans. In mice, loss of Rbfox2 in the *Nkx2-5* lineage caused embryonic lethality with hearts consisting of a common ventricle and outflow tract^18^. However, heterozygous animals were viable and phenotypically normal. We reported similar outcomes in zebrafish, where the function of RBFOX2 is split between two paralogs, *rbfox1l* and *rbfox2*^19^. Furthermore, myocardial-specific re-expression of Rbfox1l was sufficient to rescue heart structure and function in zebrafish double mutants, implicating defective myocardium as causal. While these cross-species comparisons established *RBFOX2* as an essential regulator of cardiac development, the species-specific differences in dosage sensitivity underscores the need for a human-based system to model the effects of *RBFOX2* haploinsufficiency, specifically in the cardiomyocyte (CM) lineage.

One strategy involves the use of genetically engineered human induced pluripotent stem cells (iPSCs) and their derivatives, including CMs^20^. Advantages include well-established protocols for CM differentiation and stage-specific profiling where specific cellular and molecular features have been defined along the iPSC-to-CM continuum that closely parallels fetal heart development, the time when HLHS arises^20,21^. Beyond differentiation, functional maturation can also be assessed *in vitro* using engineered heart tissues (EHTs) assembled from iPSC-CMs.

Here, we generated and characterized human iPSC-CMs and EHTs carrying heterozygous (het) or null RBFOX2 truncating alleles. Our data demonstrate that RBFOX2 is essential for progressing CM differentiation by shifting exon usage profiles, including those of *alpha-actinin-2 (ACTN2),* to more mature states. Moreover, we found that this process is facilitated by critical levels of RBFOX2 that are required for autoregulation at mutually exclusive exons encoding early and late isoforms. In het CMs, this critical threshold is never met, leading to defective autoregulation that skews the early-to-late isoform ratio and generates a protein-truncating isoform caused by exon co-inclusion. Overexpression and rescue studies in iPSC-CMs, EHTs, and zebrafish suggest that the early isoform facilitates the progenitor-to-CM transition, the late isoform promotes CM maturation, and the aberrant isoform exerts dominant-negative activity. Finally, we demonstrate that overexpression of correctly spliced ACTN2 rescues het, but not null, phenotypes by restoring contractility, which triggers a mechanosensing feedback loop involving upregulation of *RBFOX2* from the remaining wildtype allele and transcriptome maturation. Collectively, our study offers a mechanistic framework for how *RBFOX2* haploinsufficiency contributes to CHD pathogenesis and eventual HF susceptibility. By extension, correcting *RBFOX2* or *ACTN2* expression or splicing might benefit people with CHD related to impaired RBFOX2 function.

## RESULTS

### Genetically engineered human iPSCs model RBFOX2 haploinsufficiency

To model *RBFOX2* deficiencies in human cardiomyocytes (CMs), we introduced deleterious mutations in exon 3, which is upstream of the RNA Recognition Motif (RRM; Figure 1A). We isolated a heterozygous clone carrying a 1 base pair (bp) insertion in one allele (het cells) and a null clone with biallelic frameshifting mutations, including a 1 bp insertion and a 2 bp deletion (null cells). Both mutations shift the open reading frame and produce a premature stop codon. RBFOX2 deficiency did not alter stemness as iPSCs from each genotype similarly expressed the SSEA4 pluripotency marker by flow cytometry (Figure S1A)^22^. Using well-established small-molecule protocols (Figure S1B)^23,24^, control, het, and null iPSCs were differentiated to the CM lineage and efficiency was assessed by cardiac Troponin T (cTnT) staining on day 21. The percentage of cTnT-expressing cells was comparable between genotypes (Figure S1C), which also showed dose-dependent reductions in RBFOX2 transcript and protein levels (Figures 1B and S1D). These data confirm that the genetic mutations reduce RBFOX2 as expected and show that RBFOX2 deficiency does not impair iPSC pluripotency or CM identity.

**Figure 1.**
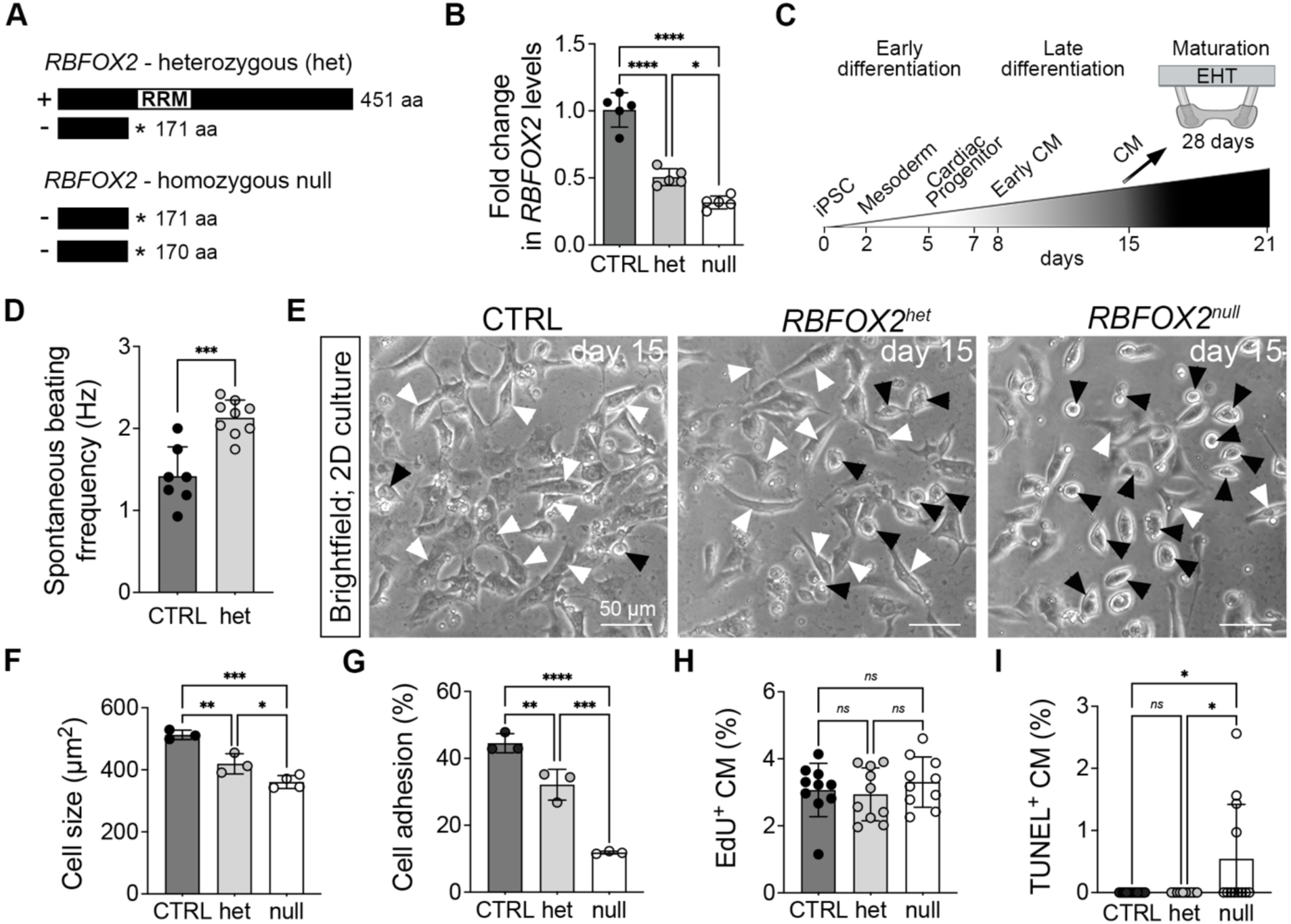
*RBFOX2* deficiency impairs cardiomyocyte growth and adhesion. (A) Schematic of predicted protein products from *RBFOX2* heterozygous (het) and homozygous null alleles. Black asterisks mark premature stop codons. RRM, RNA-recognition motif. (B) Schematic of human iPSC-directed cardiomyocyte (CM) differentiation with annotated stages: Early differentiation (days 0-7); Late differentiation (days 8-15); Maturation (3D EHT culture, days 0-28). EHT, engineered heart tissue. (C) qRT-PCR analysis of *RBFOX2* transcript levels in day 21 CTRL, het, and null iPSC-CMs (n = 5). (D) Spontaneous beating frequency of day 15 CTRL and het iPSC-CMs in 2D culture (n = 3). (E) Brightfield images of replated day 15 CTRL, het, and null iPSC-CMs after 1 day (n = 3). White arrowheads indicate flattened cells; black arrowheads indicate rounded cells. Scale bars, 50 μm. (F-I) Quantification of cell size (F; n = 3-4; each dot represents the average area of 35-55 CMs per replicate), cell adhesion (G; n = 3), percent EdU+ CM (H; n = 3; each dot represents one image count), and percent TUNEL+ CM (I; n = 3; each dot represents one image count) in day 15 CTRL, het, and null iPSC-CMs. Data are presented as mean ± SD (C, D, and F–I). *n* indicates biological replicates. Statistical analyses were performed using one-way ANOVA with Tukey’s post hoc test (C, F–I) or unpaired two-tailed *t*-test (D). Significance levels: \**p* < 0.05; \*\**p* < 0.01; \*\*\**p* **< 0.001;** \*\*\*\**p* **< 0.0001; *ns*, not significant.**

Molecular and cellular hallmarks have been defined along the directed differentiation continuum where iPSCs transition to mesoderm by day 2, CM progenitors by day 5, and early CMs by day 8 when spontaneous contractions begin. By day 15, CMs have reached a fetal-like morphological state where they are maintained in 2D culture or matured in 3D engineered heart tissues (EHTs) (Figure 1C)^20,21^. Under brightfield microscopy, we found that day 15 het CMs contract qualitatively similar to controls but with more patchy and sporadic activity (Video S1). In contrast, null CMs remained non-contractile (Video S1), mirroring the functional defects observed in *rbfox1l;rbfox2* zebrafish mutants^19^. Quantitatively, het CMs exhibited elevated spontaneous beat frequencies relative to controls (Figure 1D), which is consistent with an immature electrophysiological phenotype^25^.

During replating, we found that more het and null CMs were required to reach confluence, suggesting deficiencies in cell growth, proliferation, or survival. Indeed, a greater proportion of het and null CMs appeared smaller and rounder on day 15 than controls (Figure 1E). Quantitative analyses confirmed reduced CM areas (Figure 1F), and Vybrant adhesion assays demonstrated significantly impaired attachments to the matrix (Figure 1G). Although reduced CM proliferation has been linked to reduced left ventricular size in HLHS hearts^26^, no genotype-specific alterations in EdU incorporation were observed (Figures 1H and S1E).

Moreover, apoptotic rates assessed by TUNEL staining were also unchanged in het CMs but slightly elevated in null cells (Figures 1I and S1F). Together, these findings suggest that *RBFOX2* haploinsufficiency compromises CM growth and adhesion but not proliferation or apoptosis.

### RBFOX2 promotes CM differentiation

We next evaluated additional hallmarks of CM differentiation, including myofibril structure, gene expression-based isoform switching, electrophysiology, and mitochondrial respiration. On day 21, myofibrils and Z-discs, marked by cTNT and ACTN2, respectively, appeared less assembled in het CMs with a 2-fold increase in the myofibrillar disarray index compared to controls (Figure 2A and 2B). Null CMs expressed cTnT and ACTN2 but lacked discernible myofibrils and Z-discs altogether (Figure 2A), similar to observations in zebrafish *rbfox1l;rbfox2* mutant hearts^19^. Transmission electron micrographs further supported these observations in that Z-discs appeared less defined in het CMs and abnormal in null cells (Figure 2C). Quantifications revealed significantly reduced Z-disc heights (Figures 2D) in het CMs but no differences in Z-disc to Z-disc lengths (Figure S2A).

**Figure 2.**
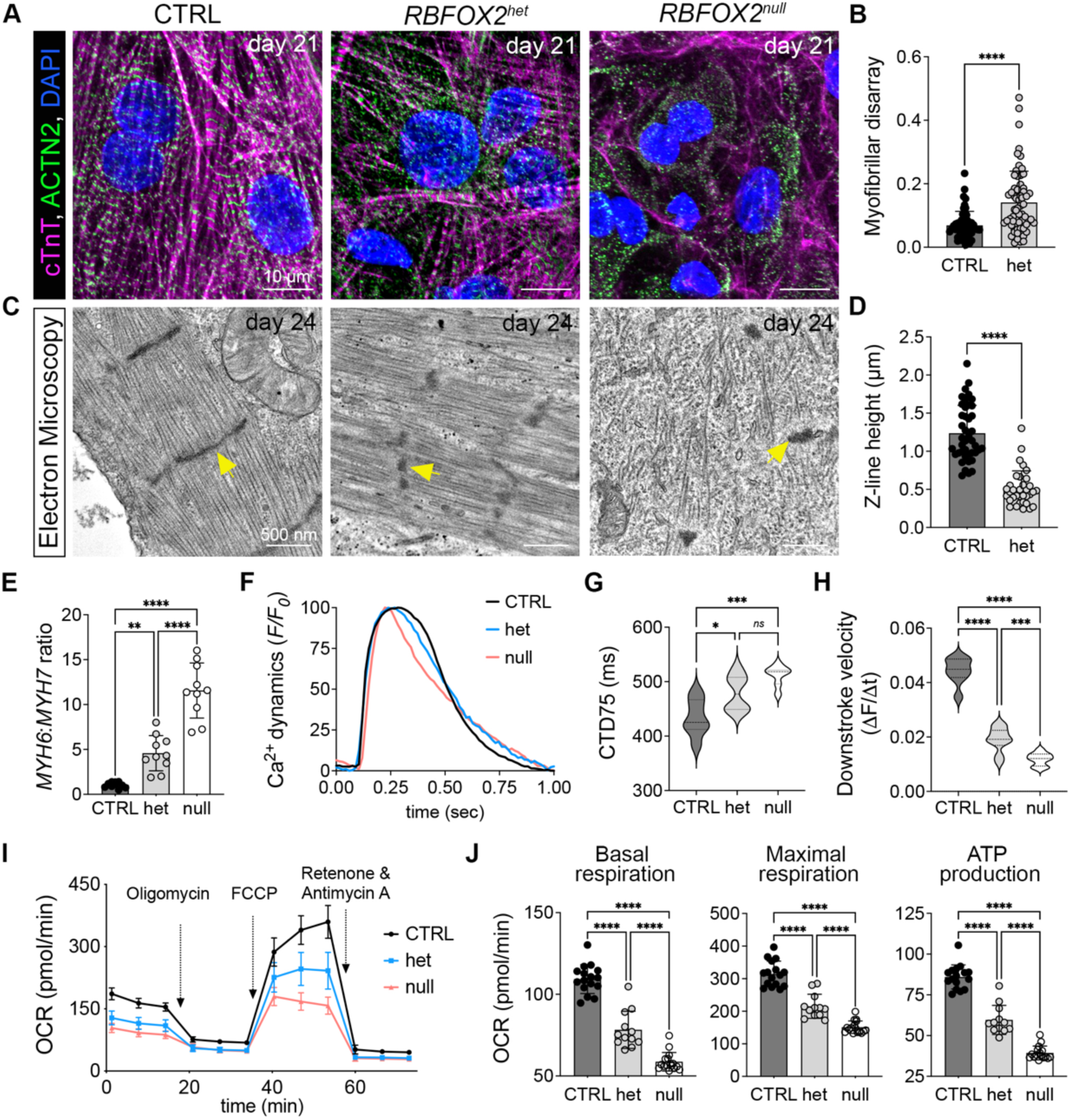
RBFOX2 promotes cardiomyocyte differentiation. (A, B) Immunofluorescence staining of day 21 CTRL, het, and null iPSC-CMs for cardiac Troponin T (cTnT, magenta), α-actinin-2 (ACTN2, green), and DAPI (blue) (A), and quantification of myofibrillar disarray (B) (n = 3; each dot represents one CM). Scale bars, 10 μm. (C, D) Transmission electron micrographs of day 24 CTRL, het, and null iPSC-CMs (C), and quantification of Z-disc heights (n = 3; each dot represents one sarcomere). Yellow arrows highlight the Z-disc (n = 3). Scale bars, 500 nm. (E) qRT-PCR analysis of *MYH6*:*MYH7* transcript ratios in day 21 CTRL, het, and null iPSC-CMs (n = 9-10). (F-H) Representative calcium transient profiles under 1 Hz pacing (F) and quantification of calcium transient duration at 75% repolarization (CTD75; G), and downstroke velocity (H) in day 21 CTRL, het, and null iPSC-CMs (n = 3). (I, J) Seahorse analysis of mitochondrial function. Profile of real-time oxygen consumption rate (OCR) in day 21 CTRL, het, and null iPSC-CMs (I), with quantification of basal respiration, ATP production, and maximal respiration (J) (n = 3). FCCP, carbonyl cyanide 4-(trifluoromethoxy)phenylhydrazone. Data are presented as mean ± SD (B, D, E, and J) or as median with interquartile range (G and H). n indicates biological replicates. Statistics are unpaired two-tailed t test (B and D) or one-way ANOVA with Tukey’s post hoc test (E, G, H and J). Significance levels: \**p* < 0.05; \*\**p* **<** 0.01; **\*\*\**p*** < 0.001; **\*\*\*\**p*** < 0.0001; *ns*, not significant.

As CM differentiate both *in vitro* and *in vivo*, their transcriptomes progressively mature through differential gene expression and alternative splicing. In terms of gene expression, immature CMs predominantly express MYH6, which is gradually replaced by MYH7 over time^20^. Accordingly, the MYH6:MYH7 ratio serves as a reliable proxy of CM differentiation status. On day 21, het and null CMs exhibited dose-dependent increases in MYH6:MYH7 ratios (Figure 2E), consistent with impaired differentiation. Calcium transients, recorded as changes in membrane potential in paced CMs, were also abnormal in both het and null cells, exhibiting prolonged durations (CTD75; Figure 2G) and slower downstroke velocities (Figure 2H). In null CMs, accelerated repolarization kinetics (Figure S3B), reflected by shortened action potential durations (APD75) were also observed (Figure S2C). These changes in electrophysiology could reflect delays in CM differentiation or a fate switch from more ventricular to atrial phenotypes. Because we did not observe decreases in ventricular marker expression or increases in atrial marker expression (Figures S2D and S2E), we conclude that these alterations are caused by stalled differentiation rather than a shift in identity.

We next assessed mitochondrial area, number, and respiration as dynamic changes have also been associated with CM differentiation and are perturbed in HLHS^27^. From transmission electron micrographs, we found no difference in mitochondrial number, but a 1.3- and 1.5-fold reduction in mitochondrial area in het and null CMs, respectively (Figures S2F and S2G). Using the Seahorse assay to measure oxygen consumption rates (OCRs), we found dose-dependent respiratory defects, including reduced basal respiration, maximal respiration, and ATP production (Figures 2I and 2J). These findings parallelled observations in *rbfox1l;rbfox2* mutant zebrafish, underscoring a conserved role for *RBFOX2* in maintaining mitochondrial function across species^19^.

Collectively, these structural, molecular, electrophysiological, and metabolic abnormalities suggest that *RBFOX2* deficiency broadly impairs CM differentiation. Notably, these phenotypes are similar to those observed in iPSC-CMs derived from HLHS patients with acute heart failure^27^, further indicating that diverse genetic lesions might converge on stalled CM differentiation.

### Molecular profiling defines expression and splicing changes in RBFOX2 deficient CMs

To define the molecular mechanisms underlying the observed phenotypes, we performed bulk RNA sequencing (RNA-seq) on control, het, and null CMs on day 21 of differentiation. Principal component analysis showed genotype-dependent clustering, indicating significant transcriptional divergence (Figure 3A). In terms of differentially expressed genes (DEGs), we found that het and null CMs retained hallmarks of an immature transcriptional profile compared to controls^28^ (Figure 3B). Specifically, het and null cells displayed dose-dependent elevations in fetal-to-adult isoform ratios in transcripts encoding structural genes, including *MYH6:MYH7*, *MYL7:MYL2*, and *TNNI1:TNNI3*. In addition, we observed dose-dependent increases in fetal ion channel expression (e.g., *HCN4*, *HCN1*, *CACNA1G*, *CACNA1H*) with reductions in the adult versions (e.g., *SCN5A*, *KCNJ2*, *CASQ2*, *PLN*).

**Figure 3.**
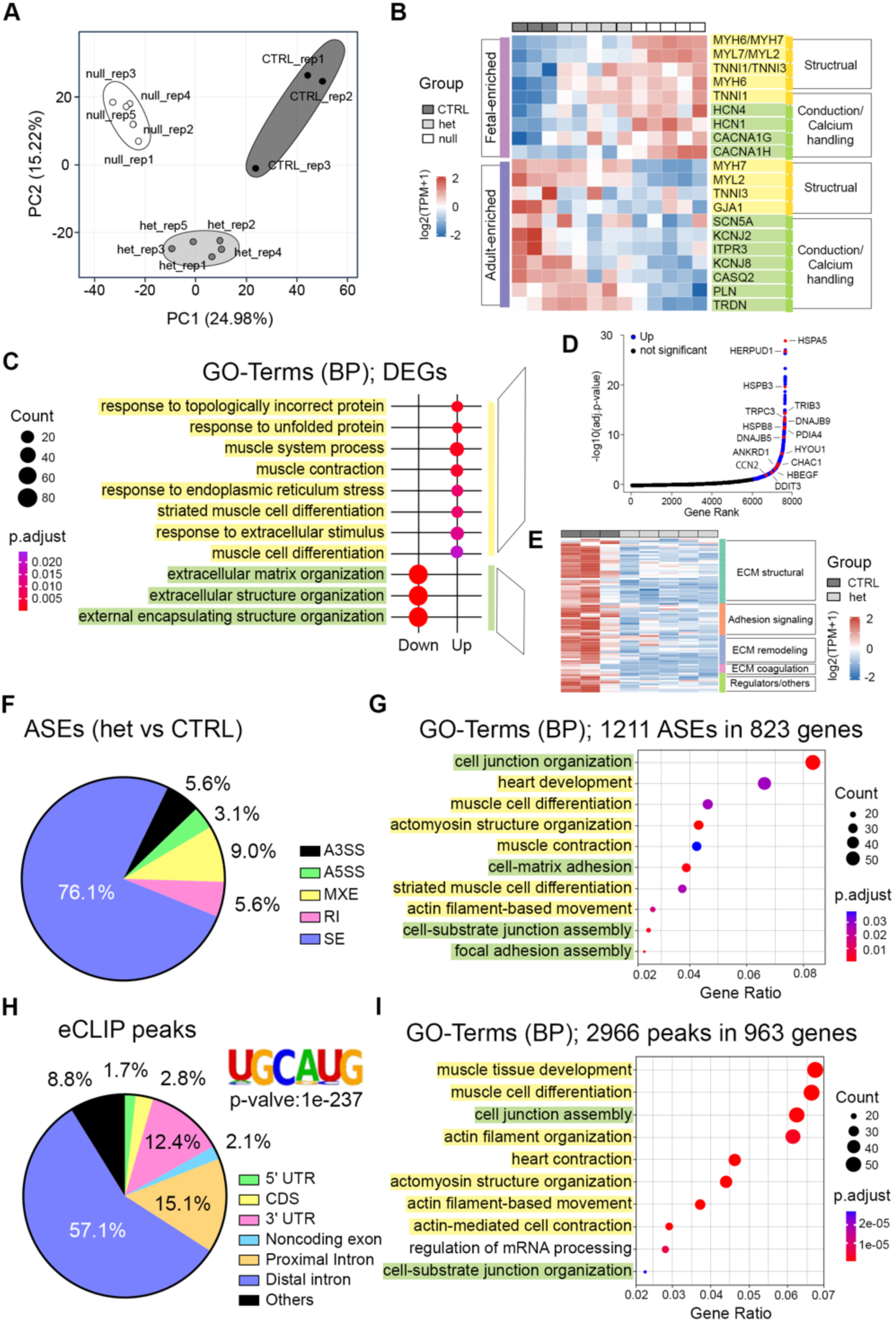
*RBFOX2* deficiency disrupts transcriptional and splicing programs. (A) Principal component analysis (PCA) of bulk RNA-seq data from CTRL (n = 3), het (n = 5), and null (n = 5) day 21 iPSC-CMs. (B) Heatmap of normalized expression from bulk RNA-seq showing immature (fetal ventricle-enriched) and mature (adult ventricle-enriched) structural, conduction, and calcium-handling transcriptional markers across CTRL (n = 3), het (n = 5), and null (n = 5) day21 iPSC-CMs. Values represent row Z-scores of log2(TPM + 1). TPM, transcripts per million. Yellow highlights structural gene or markers, and green highlights conduction, and calcium-handling genes. (C) Bubble plot of enriched GO terms associated with downregulated and upregulated differentially expressed genes (DEGs) in het iPSC-CMs compared with CTRL. BP, biological process. Yellow highlights upregulated, and green highlights downregulated GO terms. (D) Hockey stick plot of upregulated DEGs (blue) in het iPSC-CMs compared with CTRL. Representative genes (red) are highlighted (FDR < 0.05; |log2 fold change| > 0.5). (E) Heatmap of extracellular matrix (ECM)-related GO terms showing differential expression across ECM structural components (collagens, laminins, elastic fibers, proteoglycans, glycoproteins), adhesion signaling, ECM remodeling factors, ECM-coagulation pathways, and other regulators between CTRL and het iPSC-CMs (86 genes). Values represent row Z-scores of log2(TPM + 1). TPM, transcripts per million. **(F)** Distribution of alternative splicing events (ASEs) identified in het versus CTRL iPSC-CMs (FDR < 0.05; |inclusion level difference| > 0.1). **(G)** GO terms enriched among 1,211 ASEs detected in 823 genes (het vs CTRL). Yellow highlights CM differentiation-related pathways, and green highlights junction-related pathways. **(H)** Distribution of RBFOX2 eCLIP peaks across genomic features with HOMER motif analysis showing significant enrichment of the consensus UGCAUG motif (two-sided enrichment, *p <* 1e-237; log2 fold change > 3; n = 2). **(I)** GO terms enriched among 2,966 RBFOX2 eCLIP peaks in 963 genes. Yellow highlights CM differentiation-related pathways, and green highlights junction-related pathways.

Because CHD patients are haploinsufficient for *RBFOX2*, we performed Gene Ontology (GO) analysis on DEGs between het and WT samples. In the upregulated subset, we found enrichment of unfolded protein response (UPR) and endoplasmic reticulum (ER) stress terms (Figure 3C; Spreadsheet S1) that contained genes such as canonical UPR effectors (e.g., *HSPA5*(*GRP78/BiP*), *DDIT3*(*CHOP/GADD153*), *TRIB3*(*TRB3*), *CHAC1*), ER-associated degradation (ERAD) components (e.g., *HERPUD1*, *DNAJB9*), and protein-folding catalysts/chaperones (e.g., *PDIA4*, *HYOU1(ORP150)*, *HSPB8*)( Figure 3D; Spreadsheet S1)^29^. We also observed upregulation of *MANF*, which functions to protect cells from ER stress^29,30^. In addition, terms related to muscle system process were enriched that included additional stress-responsive genes such as *ANKRD1*, *CCN2*, *TRPC3*, *HBEGF*, and *DES* (Figure 3D; Spreadsheet S1). Importantly, many of these transcripts have also been identified as upregulated in failing human hearts^31–34^ and iPSC-CMs derived from HLHS patients^35^.

In the downregulated subset, we found enrichment of extracellular matrix (ECM) organization and extracellular structural assembly (Figure 3C; Spreadsheet S1) with representative genes including the major collagens (e.g., *COL1A1*, *COL1A2*, *COL3A1*, *COL5A1*, *COL5A2*, *COL6A3*), laminin subunits (e.g., *LAMA1*, *LAMA5*, *LAMB1*), integrins (e.g., *ITGA2*, *ITGA11*, *ITGB4*), ECM glycoproteins (e.g., *FN1*, *TNC*, *DCN*, *LUM*), and junctional components (e.g., *GJA1*, *S100A10*) (Figure 3E). These transcriptional changes align with the observed defects in cell adhesion (Figure 1G) and mirror previously described ECM remodeling deficiencies in *Rbfox2* null embryonic mouse hearts^18^.

Because RBFOX2 is a well-established splicing regulator, we performed replicate Multivariate Analysis of Transcript Splicing (rMATS) to identify differential alternative splicing events (ASEs). From this analysis, we discovered 1,211 ASEs across 823 genes (Figure 3F; Spreadsheet S2). The most common event was skipped exon (SE), representing 76.1% of the total. GO analysis showed a high degree of overlap with downregulated DEGs, including terms like cell junction organization and cell-matrix adhesion (Figure 3G; Spreadsheet S2). Examples of mis-spliced transcripts include *ACTN1*, *ACTN2*, *KINDLIN2*, *PTK2*, *ITGA3*, *ITGA7*, *FN1*, and *PXN*. We also identified terms associated with CM differentiation such as heart development, myofibril differentiation, and actomyosin organization with mis-spliced transcripts including *TPM1*, *TTN*, *OBSCN*, *NEBL*, *FHOD3*, *MYH10*, *PDLIM5*, and *PDLIM7* (Figure 3G, Spreadsheet S2). These data suggest that *RBFOX2*-sensitive exons are enriched in genes regulating sarcomere assembly, cytoskeletal architecture, and cell adhesion.

To determine which transcripts are directly bound by RBFOX2 in the CM lineage, we performed eCLIP-seq on day 15 WT CMs and identified 2,966 high-confidence peaks in 963 genes that were enriched for the RBFOX2 consensus binding sequence, UGCAUG (p < 1×10⁻²³⁷, Figure 3H). Consistent with previously published RBFOX2 eCLIP profiles from diverse cell types^36^, over 70% were found in introns with 57.1% in distal sequences (Figure 3H), previously defined as >500 base pairs (bp) away from an exon^36,37^. 15.1% were found in proximal introns (Figure 3H), defined as <500 bp from an exon^36^. A smaller fraction, 12.4%, were observed in 3′UTRs (Figure 3H), which can mediate transcript stability, translation, and alternative polyadenylation^37,38^. GO analysis revealed that RBFOX2-bound transcripts were significantly enriched for muscle-specific terms, such as muscle cell differentiation, heart contraction, and actin-mediated cell contraction (Figure 3I, Spreadsheet S3). Because the RBFOX2 binding profiles vary substantially between cell types and developmental stages, caution should be used when drawing conclusions based on publicly available data.

### RBFOX2 directly regulates exon usage to mature the transcriptome

To distinguish direct from indirect splicing changes, we performed dataset integration between differential ASEs identified in het CMs and eCLIP+ RBFOX2 targets. Through this analysis, we observed significant overrepresentation of GO terms including muscle structure development, cell junction organization, muscle cell differentiation, and cell junction assembly (Figure 4A; Spreadsheet S4), suggesting that RBFOX2 directly regulates these splicing programs. By projecting these changes onto an annotated map of CM-CM and CM-ECM interactions, we found alterations in genes that facilitate sarcomere assembly and force transmission during systolic contraction^39^ (Figure 4B). Specific genes affected encode sarcomere components (*TPM1*), Z-disc subunits (*PDLIM-5, ACTN2, ACTN4*), focal adhesion complex members (*ITGA7*, *KINDLIN2*), and intercellular junctional proteins (e.g., *ZO-1*).

**Figure 4.**
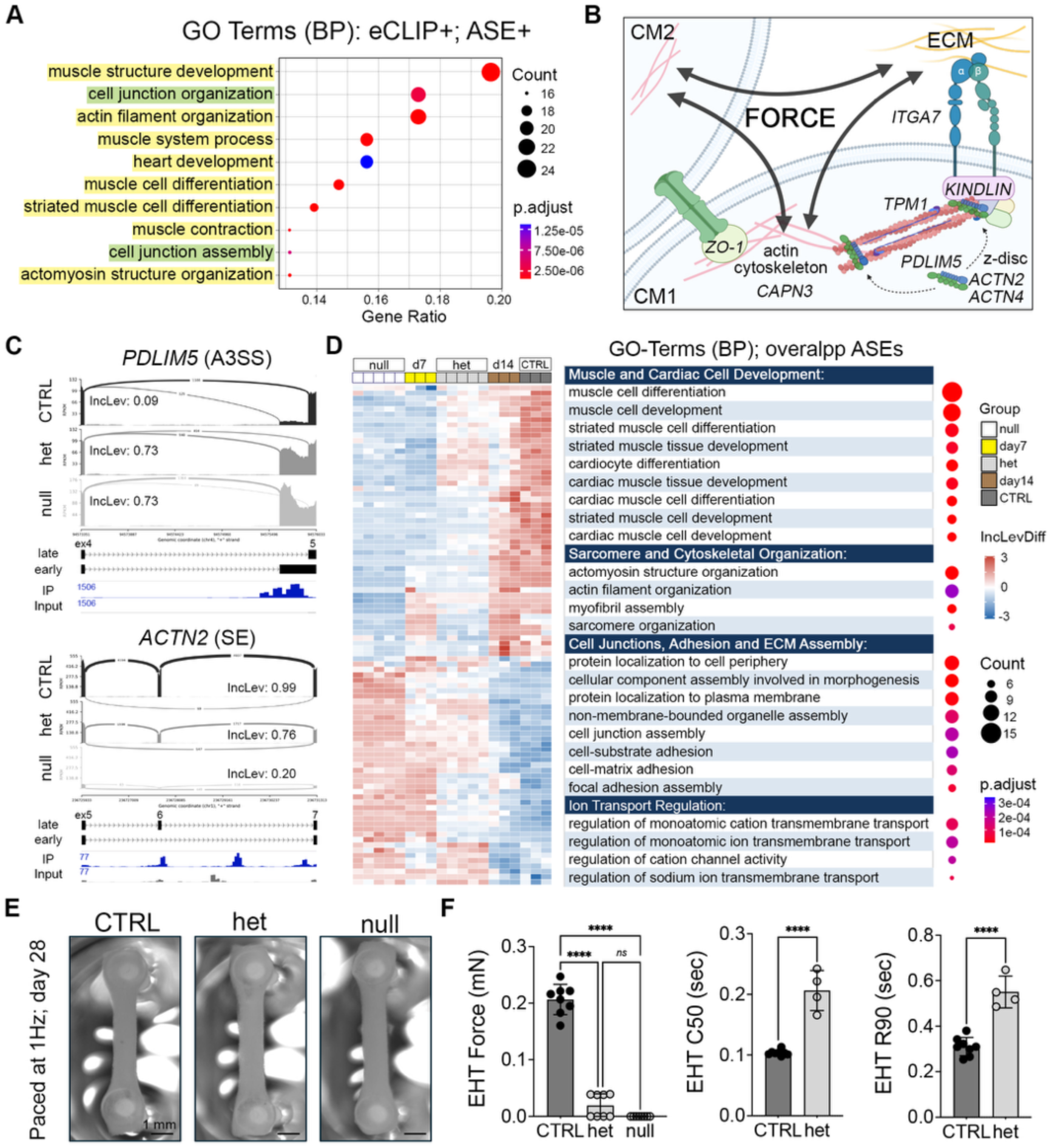
RBFOX2 shifts exon usage to a more mature profile in target transcripts encoding sarcomere, cytoskeletal, and junctional components. **(A)** Gene Ontology (GO) terms enriched among overlapping RBFOX2 eCLIP targets and alternative splicing events (ASEs) from het versus CTRL iPSC-CMs. Yellow highlights CM differentiation-related pathways, and green highlights junction-related pathways. BP, biological process. **(B)** Schematic model of *RBFOX2*-dependent regulation of sarcomere, cytoskeletal, junctional, and adhesion networks through eCLIP⁺ ASE⁺ targets that coordinate force generation and transmission. CM, cardiomyocyte; ECM, extracellular matrix. **(C)** Sashimi plots of splicing regulation at *PDLIM5* (A3SS) and *ACTN2* (SE) loci in CTRL, het, and null day 21 iPSC-CMs. Inclusion levels (IncLev) are indicated. Representative RBFOX2 eCLIP binding peaks are shown (IP) compared to input control. Numbers on tracks indicate normalized read counts. A3SS, alternative 3′ splice site; SE, skipped exon; ex, exon; early, early isoform; early, late isoform. **(D)** Heatmap of overlapping ASEs (het vs CTRL and day 7 vs day 14 CMs from a published dataset) across day 7 (n = 3), day 14 (n = 3), CTRL (n = 3), het (n = 5), and null (n = 5) day 21 iPSC-CMs. GO biological process terms enriched among ASE-associated genes are shown at right. **(E)** Brightfield images of engineered heart tissues (EHTs) generated from CTRL, het, and null iPSC-CMs at day 28 under 1 Hz pacing (n = 3). Scale bars: 1 mm. **(F)** Functional assessment of EHT contractile performance (under 1 Hz pacing, day 28), including measurements of contractile force (mN), contraction time to 50% peak (C50, sec), and relaxation time to 90% peak (R90, sec) (n = 3; each dot represents one EHT). Data in are presented as mean ± SD (F). *n* indicates biological replicates. Statistical analyses were performed using unpaired two-tailed *t*-test or one-way ANOVA with Tukey’s post hoc test (F). Significance levels: \*\*\*\**p* < 0.0001; *ns*, not significant.

**Figure 5.**
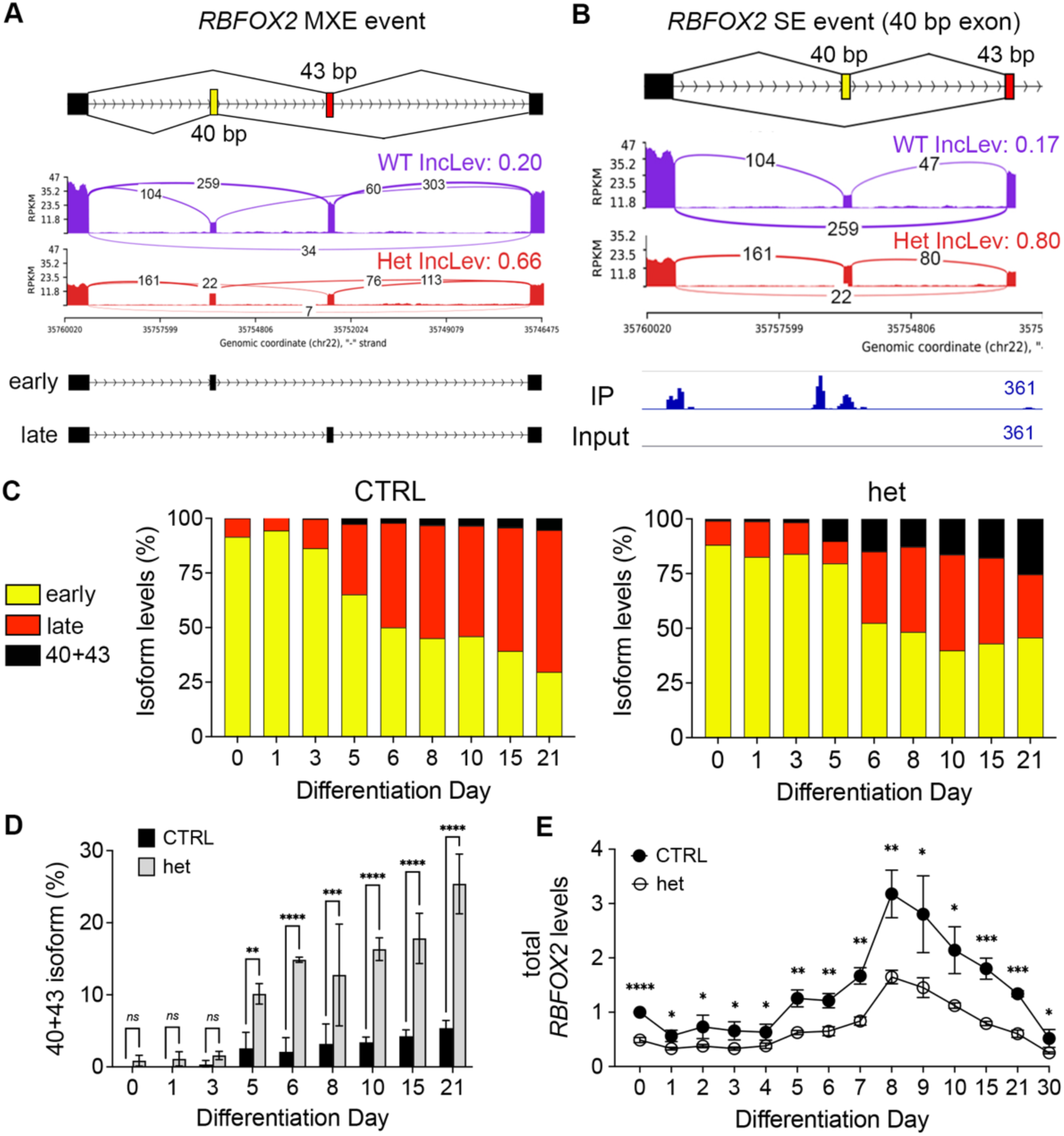
Primary defects in RBFOX2 autoregulation alters RBFOX2 isoform ratios in het CMs. (A) Schematic of *RBFOX2* mutually exclusive exons (MXEs) at the C-terminal domain, showing the early (40 bp) and late (43 bp) exons that define early versus late isoforms. Sashimi plots of *RBFOX2* exon inclusion in CTRL and het iPSC-CMs on day 21. (B) Sashimi plots of *RBFOX2* skipped exon event for the 40 bp exon in CTRL and het iPSC-CMs on day 21. Inclusion level differences (IncLev) are indicated with respect the early 40 bp exon. Plots also show aberrant co-inclusion of the 40 and 43 bp exons in WT and het iPSC-CMs. RBFOX2 eCLIP binding peaks upstream of the 40 bp exon are shown. Numbers on tracks indicate normalized read counts. SE, skipped exon. (C) DNA fragment analysis of isoform distribution of early (yellow), late (red), and aberrant 40+43 (black) *RBFOX2* transcripts in CTRL and het iPSC-CMs over a differentiation time course (days 0-21; n = 3). (D) Bar graph showing the percentage of 40+43 *RBFOX2* isoform in CTRL and het iPSC-CMs during differentiation (days 0-21; n = 3). (E) qRT-PCR analysis of total *RBFOX2* transcript levels in CTRL and het cells over a day 0-30 differentiation time course, normalized to day 0 CTRLs (n = 3). Data are presented as mean ± SD (C-E). n indicates biological replicates. Statistical analyses were performed using two-way ANOVA with Bonferroni’s post hoc test (D) or multiple unpaired *t*-tests (E). Significance levels: \**p* < 0.05; \*\**p* < 0.01; \*\*\**p* < 0.001; \*\*\*\**p* < 0.0001; *ns*, not significant.

By examining specific mis-spliced exons near RBFOX2 binding peaks, we found retention of exon usage profiles previously associated with relative immaturity^21,40^. Specifically, differential exon usage has been described between day 7 and day 14 iPSC-CMs^21^ as well as fetal and adult mouse hearts^41^. For example, we found that despite showing similar levels of *PDLIM5* expression, het cells predominantly include the longer version of exon 5 (Figure 4C), which is more frequently found in day 7 than in day 14 iPSC-CMs^21^ . In contrast, control cells almost always included the shorter exon (Figure 4C). The same exon 5 splicing defect was also observed in zebrafish and mouse *RBFOX2* null hearts^18,19^. Moreover, we found that RBFOX2 eCLIP peaks were enriched over the longer exon 5 sequence (Figure 4C), consistent with a direct role in splicing. Another example is *ACTN2* where both expression and splicing defects were observed (Figures 4C and 7A). In addition to 42% reductions in *ACTN2* expression levels, het cells more frequently skipped exon 6 (Figure 4C), which encodes the actin binding domain required for transcript stability and protein function in Z-disc and sarcomere assembly^40^. RBFOX2 eCLIP peaks were observed in the proximal intron downstream of exon 6 (Figure 4C), a position linked to exon inclusion^36^. Interestingly, we found similar fetal exon biases in *ZO-1*, *ITGA7*, *ACTN4*, *TPM1*, *CAPN3*, and *KINDLIN2* (Figures 4B and S3A), which we confirmed by qRT-PCR (Figure S3B).

**Figure 6.**
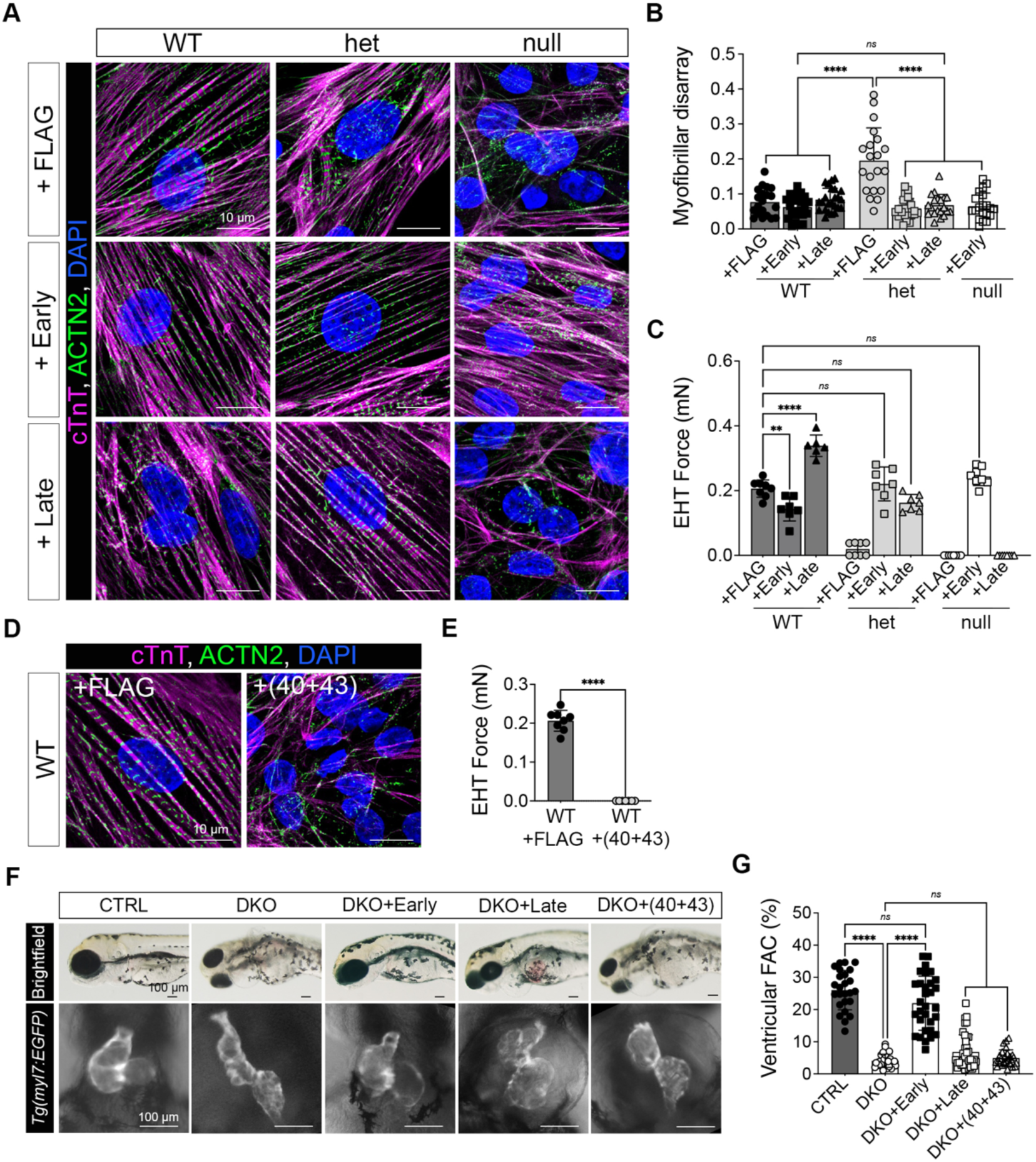
Isoform-specific roles for RBFOX2 in CM differentiation and maturation. (A, B) Immunofluorescence staining of day 15 WT, het, and null iPSC-CMs transduced with 3×FLAG control (+FLAG), Early, or Late *RBFOX2* isoforms, labeled with cTnT (magenta), labeled with ACTN2 (green), and DAPI (blue) (A), and quantification of myofibrillar disarray (B) (n = 3; each dot represents one CM). Scale bars, 10 μm. (C) Quantification of EHT contractile force (under 1 Hz pacing, day 28) in WT, het, and null iPSC-CMs expressing FLAG, Early, or Late isoforms (n = 3; each dot represents one EHT). (D) Immunofluorescence staining of WT iPSC-CMs transduced with FLAG control or aberrant 40+43 isoform, labeled with cTnT (magenta), ACTN2 (green), and DAPI (blue) (n = 3). Scale bar, 10 μm. (E) Quantification of EHT contractile force (under 1 Hz pacing, day 28) in WT iPSC-CMs transduced with FLAG control or aberrant 40+43 isoform (n = 3; each dot represents one EHT). (F, G) Brightfield images of 72 hpf zebrafish embryos and *Tg(myl7:GFP)* fluorescent overlays of the heart in CTRL, DKO, and DKO embryos injected with Early, Late, and 40+43 *RBFOX2* isoform mRNA (F), and quantification of zebrafish ventricular fractional area change (FAC%, G). Scale bar, 100 μm. Data are presented as mean ± SD (B, C, E and G). n indicates biological replicates. Statistical analyses were performed using one-way ANOVA with Tukey’s post hoc test (B, C, and G) or unpaired two-tailed *t*-test (E). Significance levels: *\*\*p* < 0.001; *\*\*\*\*p* < 0.0001; *ns*, not significant.

**Figure 7.**
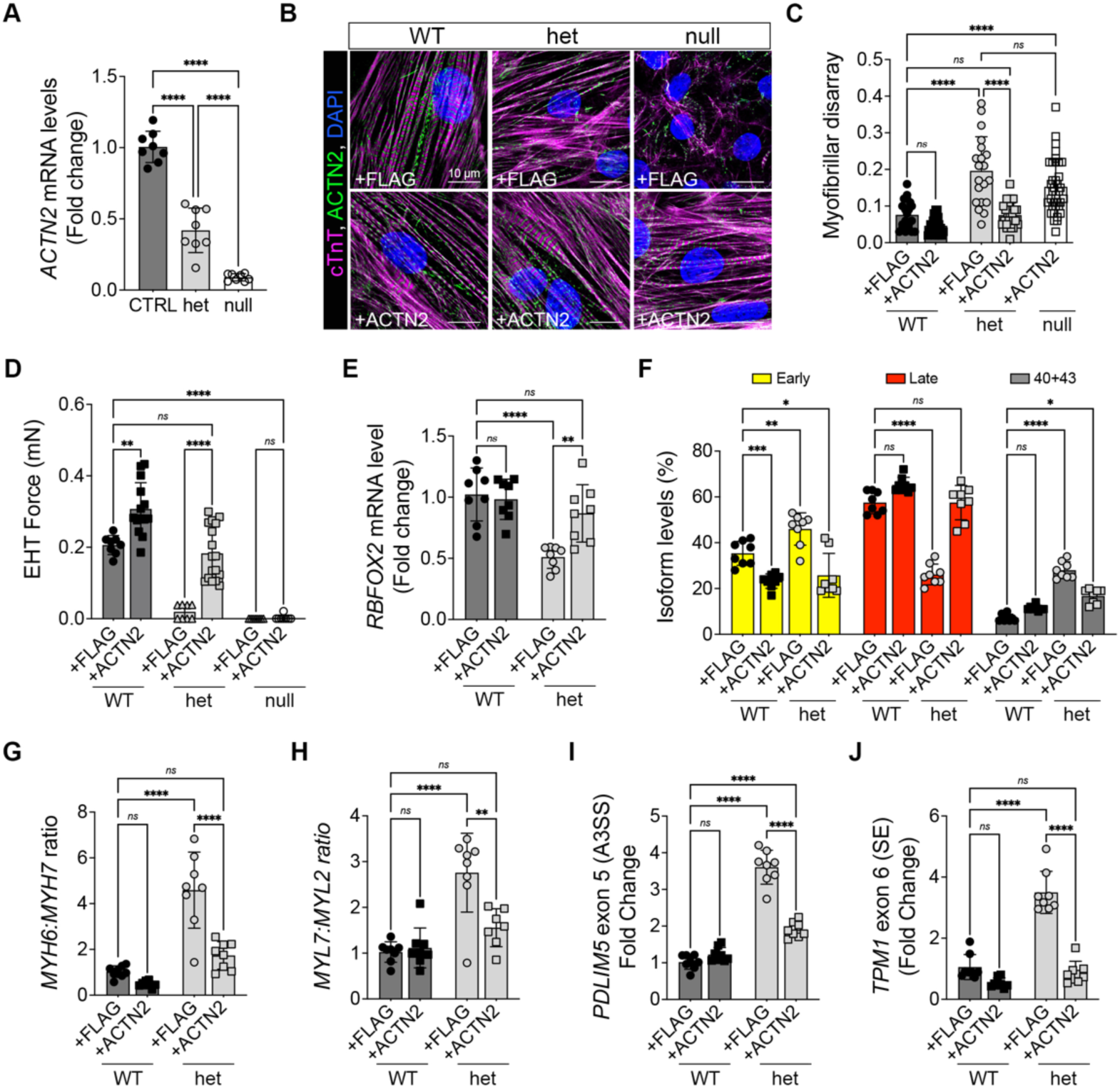
ACTN2 restores contractility in het CMs and triggers a mechanosensitive feedback loop that reactivates RBFOX2 expression and transcriptome maturation. (A) qRT-PCR analysis of *ACTN2* transcript levels in day 21 CTRL, het, and null iPSC-CMs (n = 8). (B, C) Immunofluorescence staining of day 15 WT, het, and null iPSC-CMs transduced with 3×FLAG control (+FLAG) or ACTN2 late isoform (+ACTN2), labeled with cTnT (magenta), ACTN2 (green), and DAPI (blue) (B), and quantification of myofibrillar disarray (C) (n = 3; each dot represents one CM). Scale bar, 10 μm. (D) Quantification of EHT contractile force (under 1 Hz pacing, day 28) in WT, het, and null iPSC-CMs transduced with FLAG or ACTN2 late isoform (n = 3; each dot represents one EHT). (E) qRT-PCR analysis of *RBFOX2* transcript levels in day 21 WT and het iPSC-CMs transduced with FLAG or ACTN2 late isoform (n = 8). (F) DNA fragment analysis of isoform percentages of early, late, and aberrant 40+43 *RBFOX2* transcripts in day 21 WT and het iPSC-CMs transduced with FLAG or ACTN2 late isoform (n = 8). (G, H) qRT-PCR analysis of *MYH6*:*MYH7* transcript ratio (G) and *MYL7*:*MYL2* transcript ratio (H) in day 21 WT and het iPSC-CMs transduced with FLAG or ACTN2 late isoform (n = 7-8). (I, J) qRT-PCR analysis of alternative splicing events *PDLIM5* exon 5 (A3SS), and *TPM1* exon 6 (SE). Isoform expression levels were normalized to total transcript levels in day 21 WT and het iPSC-CMs transduced with FLAG or ACTN2 late isoform (n = 7-8). A3SS, alternative 3′ splice site; SE, skipped exon. Data are presented as mean ± SD (A, and C-J). *n* indicates biological replicates. Statistical analyses were performed using one-way ANOVA with Tukey’s post hoc test. Significance levels: \**p* < 0.05; \*\**p* < 0.01; \*\*\**p* < 0.001; \*\*\*\**p* < 0.0001; *ns*, not significant.

Based on these observations, we hypothesized that RBFOX2 might facilitate more global transcriptome maturation events through alternative splicing. To test this idea, we integrated our datasets from day 21 control, het, and null CMs with publicly available day 7 and day 14 datasets^21^ from the same parental strain. We found 95 ASEs in 68 independent genes that were enriched for GO terms involving muscle and cardiac cell development, sarcomere and cytoskeletal organization, cell junctions, adhesion and ECM assembly, and ion transport regulation (Figure 4D; Spreadsheet S5). Most strikingly, heatmaps revealed that our day 21 control CMs displayed more advanced splicing profiles than those observed on day 14, indicative of increased transcriptome maturation (Figure 4D). In contrast, *RBFOX2*-deficient cells failed to progress along this continuum with null CMs exhibiting less advanced profiles than those on day 7, and het CMs showing an intermediate phenotype (Figure 4D). Overall, these splicing abnormalities are consistent with dose-dependent requirements for RBFOX2 in promoting global transcriptome maturation, which supports CM differentiation.

Based on these molecular profiles, we suspected that contractile function might be more significantly impaired in the context of EHTs^42^ than in monolayer culture as the matrix environment, which promotes cell-cell and cell-ECM attachments, is likely compromised^43^. We generated EHTs from WT, het, and null CMs using standard methods^44^, cultured them for 28 days, and quantified metrics of contractility in paced samples. Unlike wild type (WT) EHTs that showed robust force with typical contraction-relaxation kinetics, null EHTs were non-contractile (Figures 4E and 4 F, Video S2), reflecting their 2D phenotype (Video S1). However, het EHTs appeared more functionally compromised than their 2D counterparts with significant reductions in force generation and prolonged contraction-relaxation times compared to controls (Figures 4E and 4F; Video S2). Collectively, these data demonstrate that RBFOX2 deficiency disrupts transcriptome maturation, CM differentiation, and function at the tissue-level.

### Primary defects in RBFOX2 autoregulation alter isoform ratios in het CMs

We next sought to identify initiating molecular events that might drive the widespread mis-splicing events observed in het CMs. Among the eCLIP+ transcripts with differential ASEs was *RBFOX2* itself, which was mis-spliced at previously described mutually exclusive exons, called B40 and M43, encoding distinct C-terminal domains (Figure 5A)^41,45,46^. Inclusion of the 40 bp exon (B40) produces an amino acid sequence containing three tyrosine residues that promotes assembly of the large assembly of splicing regulators (LASR), a higher-order protein complex that enhances splicing activity^45^. In contrast, inclusion of the 43 bp exon (M43) generates a sequence lacking these tyrosines, which prevents LASR formation. Exon inclusion was also shown to be developmentally regulated^41^. Specifically, mouse and human CMs predominantly included the 40 bp exon during fetal life, which became almost completely replaced by the 43 bp exon in adulthood^41^.

Consistent with stalled differentiation, het CMs preferentially included the 40 bp early exon on day 21, while controls favored the 43 bp late exon (Figure 5A and 5B). When looking at the 40 bp skipped exon event specifically, we also noticed that both WT and het CMs produced an aberrant isoform that included both exons (40+43; Figure 5B), which shifts the open reading frame, introduces a premature stop codon, and truncates the protein. This truncation alters the C-terminal splicing domain while preserving the RNA Recognition Motif (RRM), suggesting it might have dominant negative activity. Consistent with autoregulation, RBFOX2 eCLIP peaks were also enriched in the proximal intron upstream of the 40 bp early exon (Figure 5B), a position associated with exon exclusion^45^. From these data, we conclude that exclusion of the early exon is sensitive to RBFOX2 dosage, which when reduced compromises transcript integrity once the late exon starts to be added.

To define the ratio of isoforms produced from the MXE event, we performed fluorescent DNA fragment analysis over a developmental time course (Figure 5C). Control cells showed progressive declines in early isoform expression beginning on day 5 and continuing through day 21 with reciprocal increases in the late isoform, consistent with a developmental shift in *RBFOX2* isoform expression. While this transition was significantly comprised in het CMs, the most profound difference was the continual rise in 40+43 isoforms (Figures 5C and 5D).

Compared to WT CMs that expressed 2.6% and 5.4% on days 5 and 21, respectively, het CMs expressed 10.1% and 25.4%, representing a 3.9-4.7-fold increase over time. Moreover, the appearance of 40+43 isoforms coincided with *RBFOX2* upregulation on day 5, which peaked on day 8 before subsequently declining. Although similar patterns in *RBFOX2* expression were observed in het CMs, levels were consistently reduced by half (Figure 5E).

Overall, our data support a model where *RBFOX2* expression is upregulated on day 5, reaching the level required to exclude the early 40 bp exon. In het CMs, this critical threshold is never met, resulting in sustained early exon inclusion. This initial defect becomes problematic when the late exon starts to be added, resulting in progressive accumulation of 40+43 transcripts with potential dominant-negative activity. Although these abnormal transcripts also exist in WT CMs, they are 4 to 5-fold less abundant than in het cells, where they constituted 25% of total *RBFOX2* mRNA. Thus, the primary defect caused by *RBFOX2* haploinsufficiency is inadequate levels of RBFOX2 to consistently repress early exon inclusion, which leads to accumulation of 40+43 isoforms with potential dominant negative effects.

### Functional assessments of RBFOX2 isoforms

To define the functions of each RBFOX2 isoform, we performed overexpression and rescue studies in WT, het, and null CMs. iPSCs were transduced with lentivirus to stably overexpress epitope-tagged early, late, and 40+43 isoforms or the epitope tag alone (+FLAG). Immunoblot analysis confirmed robust RBFOX2 isoform overexpression compared to baseline (Figure S6A).

Focusing on the early and late isoforms, we performed directed differentiation of iPSC clones from each group and assessed sarcomere assembly and organization on day 21 by immunofluorescence (Figures 6A and 6B). Overexpression of either early or late isoforms in WT CMs had no effect, suggesting high levels of either are non-toxic in this context. Compared with the +FLAG control, overexpression of the early isoform was sufficient to rescue the myofiber disarray index in both het and null CMs, whereas the late isoform was only effective in het cells. These findings suggest that the early isoform is essential to support the initial stages of CM differentiation, which cannot be compensated for by the late isoform alone.

In assessments of contractile performance, overexpression of the early isoform in WT EHTs reduced force generation by 1.44-fold, while overexpression of the late isoform augmented it by 1.64-fold. These changes were accompanied by the predicted alterations in contraction and relaxation times (Figures 6C and S5A). In het EHTs, both isoforms restored contractile force to control levels and normalized contraction–relaxation kinetics (Figures 6C and S5A). However, in null EHTs, only the early isoform was sufficient for rescue (Figures 6C and S5A). Consistent with our findings from 2D culture, these observations support distinct requirements for each RBFOX2 isoform, with the early version supporting CM differentiation and the later version promoting maturation.

Because 40+43 isoforms are predicted to have dominant-negative activity, we tested this directly through overexpression in WT iPSCs (Figure S4B). As described above, we evaluated sarcomere assembly by immunofluorescence on day 21 and contractility in EHTs on day 28. Strikingly, CMs overexpressing 40+43 isoforms failed to form sarcomeres altogether, phenocopying the sarcomerogenesis defects observed in null CMs (Figure 6D). Similarly, EHTs were non-contractile (Figure 6E), demonstrating potent dominant-negative functions.

To determine whether these isoform-specific roles are conserved *in vivo*, we performed rescue studies in zebrafish *rbfox*1l;*rbfox2* double knock-out embryos (DKO) by injecting mRNA encoding early, late, or 40+43 human RBFOX2 isoforms. Consistent with our previous findings, early isoform overexpression largely suppressed pericardial edema and restored ventricular morphology in mutant animals (Figure 6F)^19^. By contrast, late or aberrant isoform overexpression failed to rescue either phenotype (Figure 6F). Quantification of fractional area change (%FAC) confirmed that only the early RBFOX2 isoform, but not the late or 40+43 versions, restored contractility in DKO hearts (Figure 6G). Together, these findings support a division of labor between the early and late RBFOX2 isoforms during CM differentiation and functional maturation, respectively, and suggest that the 40+43 isoform is pathogenic.

### ACTN2 overexpression rescues RBFOX2 haploinsufficiency through a positive feedback loop

*RBFOX2* deficiency disrupts the expression and splicing of genes connecting the sarcomere to the ECM, including *ACTN2* (Figures 4B and 4C)^47^, which has been previously implicated in sarcomerogenesis, CM maturation, and force transmission^48–50^. To perform these functions, ACTN2 requires its actin-binding domain (ABD), encoded by exon 6^40^. As previously shown for another RNA binding protein, RBM24^40^, we found that exon 6 inclusion also requires RBFOX2, which binds downstream of the exon in a position known to promote inclusion (Figure 4C)^36^. Furthermore, addition of exon 6 begins on day 5 of CM differentiation^40^, coinciding with *RBFOX2* upregulation (Figure 5E). Importantly, *ACTN2* transcripts lacking exon 6 are highly unstable, likely due to a frameshift that introduces a premature stop codon and triggers non-sense mediated decay (NMD)^40^. Consistent with this mechanism, qPCR analysis confirmed dose-dependent reductions in *ACTN2* mRNA levels in het and null CMs (Figure 7A). Together, these findings suggest that overexpression of exon 6-containing ACTN2 in het and null CMs might rescue sarcomere assembly, sarcomere organization, and contractility in 2D culture and EHTs.

We transduced WT, het, and null iPSCs with lentivirus to stably overexpress epitope-tagged exon 6-containing ACTN2 (+ACTN2) or the epitope tag alone (+FLAG)^48,49^. Immunoblot analysis confirmed robust ACTN2 overexpression compared to baseline (Figure S4C). In both het and null CMs, ACTN2 improved sarcomere assembly and organization on day 21 compared to the FLAG tag alone (Figure 7B). Quantifications of the myofibrillar disarray index confirmed rescue to WT levels in het CMs (Figure 7C). However, rescue was only partial in null CMs, reaching levels comparable to het CMs + FLAG. In line with these findings, contractility was restored by ACTN2 in het CMs but showed only partial improvement in nulls.

In WT EHTs, ACTN2 overexpression augmented contractile force and contraction-relaxation kinetics, indicative of increased maturation (Figures 7D and S5B). Consistent with our findings in 2D culture, ACTN2 overexpression rescued contractile force and contraction-relaxation times in het EHTs, but not null. Remarkably, these findings demonstrate that overexpression of ACTN2 alone can bypass RBFOX2 haploinsufficiency, but not full deletion, to restore CM differentiation and function. Moreover, this discrepancy between genotypes suggest that Z-disc stabilization and contractile function might initiate a positive feedback loop that matures the transcriptome through RBFOX2 upregulation, which would only be possible in the presence of a WT allele.

Indeed, ACTN2 restored total *RBFOX2* expression to WT levels in het CMs (Figure 7E). We also found restoration of more appropriate isoform ratios in het cells with lower early, higher late, and decreased 40+43 *RBFOX2* transcripts (Figure 7F). Because *RBFOX2* levels and isoform ratios were rescued, we assessed whether the more global RBFOX2-dependent maturation programs might also be restored. Indeed, fetal-to-adult isoform ratios for *MYH6:MYH7* and *MYL7:MYL2* were rescued back to WT levels in het CMs (Figures 7G and 7H) and exon usage for *PDLIM5* and *TPM1* were also restored (Figures 7I and 7J). These results demonstrate that ACTN2 overexpression can bypass *RBFOX2* haploinsufficiency to rescue CM differentiation and function by initiating a mechanosensing feedback loop that matures the transcriptome through *RBFOX2* upregulation.

## DISCUSSION

Here, we present a human model of *RBFOX2* haploinsufficiency and identify molecular defects in *RBFOX2* expression and autoregulation as initiating events that disrupt transcriptome maturation, which compromises CM differentiation and adhesion. These cellular defects were so severe that het and null CMs could not survive standard freeze-thaw protocols or assemble into embryoid bodies (EBs) under typical bioreactor conditions. Similar CM differentiation and adhesion defects that ultimately impair heart development have also been reported in null mice when *RBFOX2* is deleted at the CM progenitor stage (*Nkx2-5* lineage)^18^. By contrast, deletion of *RBFOX2* after CM differentiation has occurred (*Mlc2v* lineage) causes no developmental abnormalities^51^. Together, these data suggest that RBFOX2 is essential during the progenitor-to-CM transition, but dispensable for subsequent steps of cardiac morphogenesis once a critical stage of differentiation is reached.

Defects in CM differentiation that are also associated with retention of immature molecular states have been increasingly recognized as a common phenotype among genetically heterogeneous HLHS cases^35,52,53^. Specifically, molecular profiling of infant HLHS myocardium revealed retention of fetal-like transcriptional programs^35^. Likewise, single cell RNA-sequencing demonstrated distinct clustering of day 14 iPSC-CMs derived from control and HLHS individuals, with the latter enriched in progenitor populations and depleted from more mature ones^35^. Similar findings have been reported in iPSC-CMs derived from independent HLHS probands^27,52^. Because impaired CM differentiation has been consistently observed across genotypes in both 2D and 3D *in vitro* systems, these data collectively suggest that HLHS is not merely a consequence of altered hemodynamics but from defects intrinsic to the myocardium.

We also found that contractility is sensitive to RBFOX2 dosage, mirroring the functional abnormalities observed in iPSC-CMs derived from HLHS patients with diverse genetic etiologies^27,35,52^. On a molecular level, our RNA-seq analysis showed upregulation of UPR, ERAD, and fetal gene programs, consistent with findings in HLHS patient-derived iPSC-CMs^27,35^ and adult HF samples^54–58^. In mouse models, *RBFOX2* expression was found to be decreased in the context of pressure overload-induced heart failure, which was also associated with retention of target fetal exons^51,59^. Moreover, CM-specific deletion of *RBFOX2* led to dilated cardiomyopathy followed by HF and lethality in mice^51^, providing *in vivo* relevance to the functional deficiencies observed *in vitro*. Although HF outcomes in patients with HLHS caused by *RBFOX2* haploinsufficiency remain unknown, these collective data suggest that these individuals may be at elevated risk for HF and likely express more fetal-like transcriptomes.

Particularly intriguing are the potentially distinct molecular mechanisms underlying stalled CM differentiation and transcriptome maturation in *RBFOX2* null versus het CMs. In null CMs, exon usage fails to shift toward more mature patterns due to complete absence of RBFOX2 protein. In contrast, het CMs present a more complex scenario in which overall expression is reduced by half, and isoform ratios, derived from the wildtype allele, are skewed with a substantial fraction encoding a truncated protein with dominant negative activity. As a result, RBFOX2 activity is likely lowered by more than half of that found in controls.

Comparable defects resulting from altered RBFOX2 levels and isoform production have also been described in other disease contexts. For example, diabetic patients frequently develop cardiovascular complications, including cardiomyopathy^60^. In both a mouse model of type 1 diabetes and in patients with type 2 diabetes, elevated glucose levels increase *RBFOX2* expression in the heart, which promotes skipping of exon 6 in the *RBFOX2* transcript^61^. Exclusion of this exon, which encodes half the RRM, generates a dominant negative product that causes mis-splicing of RBFOX2 target RNAs prior to the onset of disease phenotypes^46,61–63^, suggesting a causal role.

Similarly, patients with myotonic dystrophy type 1 (DM-1) often develop cardiac conduction delays and ventricular arrhythmias in adulthood^64–66^. This cardiac phenotype has been linked to aberrant overexpression of the early RBFOX2 isoform in both DM-1 patient hearts and a corresponding mouse model^41^. Mice engineered to overexpress or exclusively express wildtype levels of the early RBFOX2 isoform live to adulthood but develop cardiac conduction defects that recapitulate the human disease phenotypes. These phenotypes are consistent our findings, which suggest distinct roles for the early and late RBFOX2 isoforms in CM differentiation and maturation, respectively.

In the context of CHD, determining whether altered RBFOX2 expression levels, isoform imbalance, or dominant negative effects contributes most to the observed phenotypes will require more rigorous genetic testing. For example, it will be informative to assess how CM differentiation and contractility are affected in iPSCs engineered to exclusively express wildtype levels of either the early or late isoform, or to express half-doses of the early isoform in the absence of the dominant negative protein. Even without such data, the identification of variants in four HLHS probands predicted to truncate RBFOX2 in the same region as the dominant negative isoform supports the idea that this protein product is pathogenic^15,16^. It will also be important to determine whether CMs heterozygous for missense *RBFOX2* variants, which potentially decrease or alter RBFOX2 activity, also accumulate 40+43 isoforms or have different molecular properties that undermine RBFOX2 function.

One mechanism by which RBFOX2 haploinsufficiency can be overcome is through overexpression of correctly spliced ACTN2, which is required for sarcomerogenesis and CM maturation^48–50^. Similar findings have been reported for another cardiac-enriched RNA binding protein, RBM24^40^, providing precedence for this mechanism. Specifically, RBM24 null human embryonic stem cell (hESC)-derived CMs display defects in CM differentiation and *ACTN2* splicing that mirror those observed in *RBFOX2*-deficient cells. In both contexts, ACTN2 overexpression rescues these cellular phenotypes, suggesting that RBM24 and RBFOX2 cooperate to promote sarcomerogenesis through *ACTN2* splicing regulation.

Conceptually, enhanced contractility is likely the result of restored sarcomerogeneis, which leads to improved Z-disc integrity and more efficient force transmission across the myofibrillar lattice. Similar findings of improved function have been documented following overexpression of mature cardiac troponin I (TNNI3) isoforms^67^. Moreover, we find that functional improvement following ACTN2 overexpression in *RBFOX2* het CMs triggers a mechanosensing feedback loop that upregulates *RBFOX2* expression, thereby normalizing its autoregulation and splicing patterns. However, in the absence of RBFOX2, this feedback mechanism cannot function, explaining why contractile force is not recovered in null EHTs. Interestingly, we also observed increased force generation in WT EHTs overexpressing ACTN2, without a corresponding rise in *RBFOX2* levels, indicating that tissue maturation can also occur through RBFOX2-independent mechanisms. Whether rescue in het CMs is specific to ACTN2 or can be replicated by other Z-disc components, such as PDLIM-5, remains to be determined.

In summary, our study provides a mechanistic framework for how RBFOX2 haploinsufficiency contributes to CHD. By extension, our findings suggest potential therapeutic strategies might involve overexpressing RBFOX2 or ACTN2 or using anti-sense oligonucleotides (ASOs) to correct *RBFOX2* splicing. While such interventions are unlikely to be administered early enough to prevent CHD *in utero*, their delivery after birth could help prevent ventricular dysfunction that often progresses to HF.

## Supporting information

Manuscript For bioRxiv_v2

## RESOURCE AVAILABILITY

### Lead contact

Further information and requests for resources and reagents should be directed to and will be fulfilled by the lead contact, Caroline E. Burns and C. Geoffrey Burns

## Materials availability

Requests for resources and reagents are available from the lead contact upon request.

## Data and code availability

- The Bulk RNA-seq and eCLIP-seq datasets generated and/or analyzed during this study have been deposited in the Gene Expression Omnibus (GEO) under accession numbers GEO: XXXX and GEO: XXXXX, respectively, and are publicly available as of the date of publication.
- This paper does not report original code.
- Any additional information required to reanalyze the data reported in this paper is available from the lead contact upon request.

## ACKNOWLEDGMENTS

We thank the Boston Children’s Hospital Aquatics Facility for zebrafish care. Sanger sequencing and DNA fragment analysis were performed at the Massachusetts General Hospital (MGH) Computational and Integrative Biology DNA Core. Transmission Electron Microscope imaging was performed at the Harvard Medical School (HMS) Electron Microscopy Core. We thank Christopher S. Chen (Boston University) for providing the pRRL-mApple–α-actinin-2 plasmid.

## AUTHOR CONTRIBUTIONS

M.H. designed, performed, and interpreted the experiments. F.L. assisted with lentivirus production, rescue experiments, data quantification, and interpretations regarding ACTN2 overexpression. Y.W., R.Z., and K.C. performed bioinformatics analyses and dataset integration. M.P., Y.T., and W.T.P. provided training to M.H. for iPSC and EHT work. J.M. provided code for EHT contractility analysis. M.P. and A.A.A. prepared eCLIP samples from 3D bioreactor WT cardiomyocytes and S.A., B.A.Y., and G.W.Y. performed eCLIP-seq experiments and bioinformatic analyses. M.T., H.Y., and V.J.B. contributed to data analysis and discussion. S.U.M. provided information on RBFOX2 variants prior to publication. C.G.B. and C.E.B. co-directed the study, secured funding, designed experiments, and interpreted data. M.H. generated the figures. M.H. and C.E.B. wrote the manuscript with input from all authors.

## KEY RESOURCES TABLE

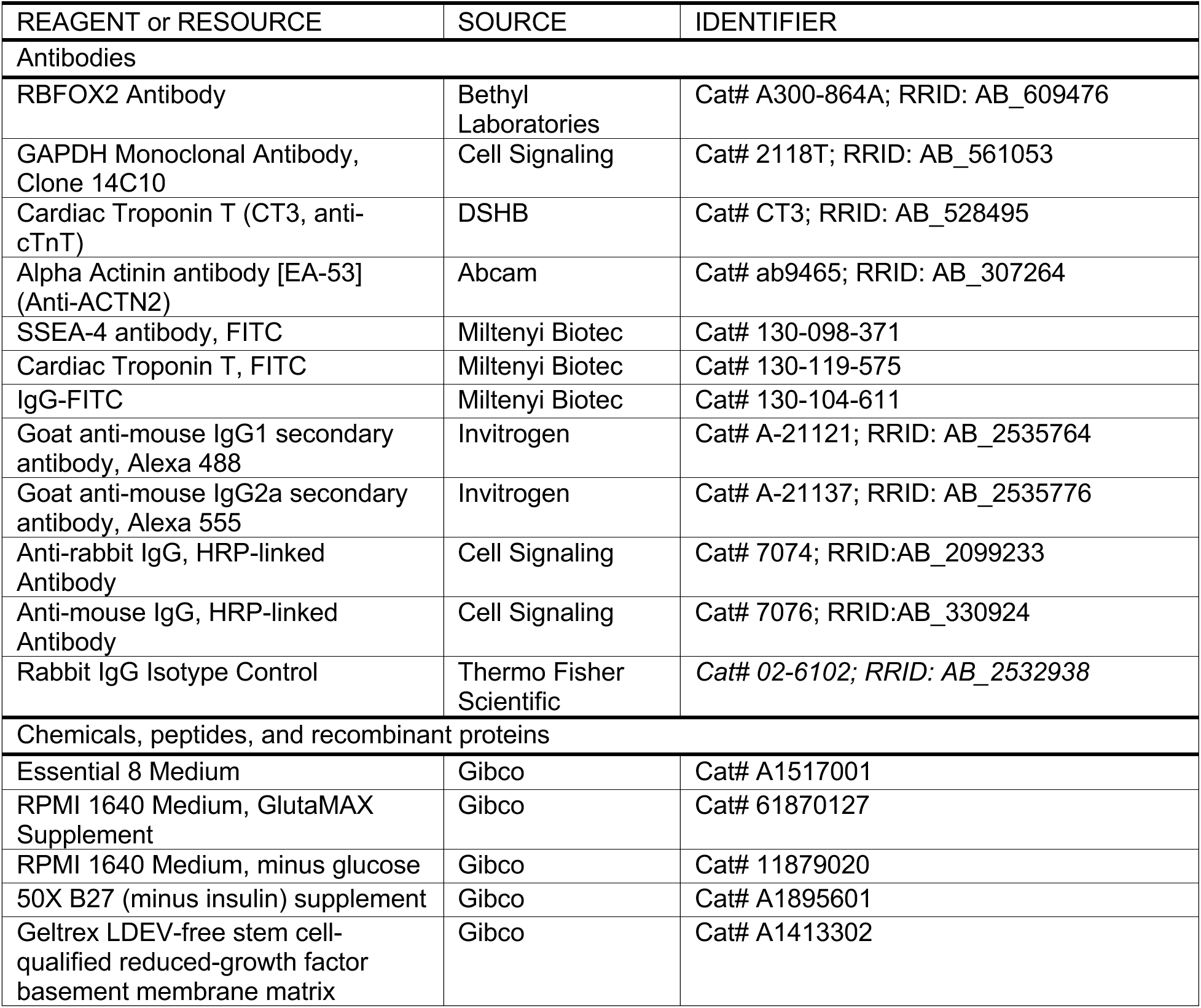
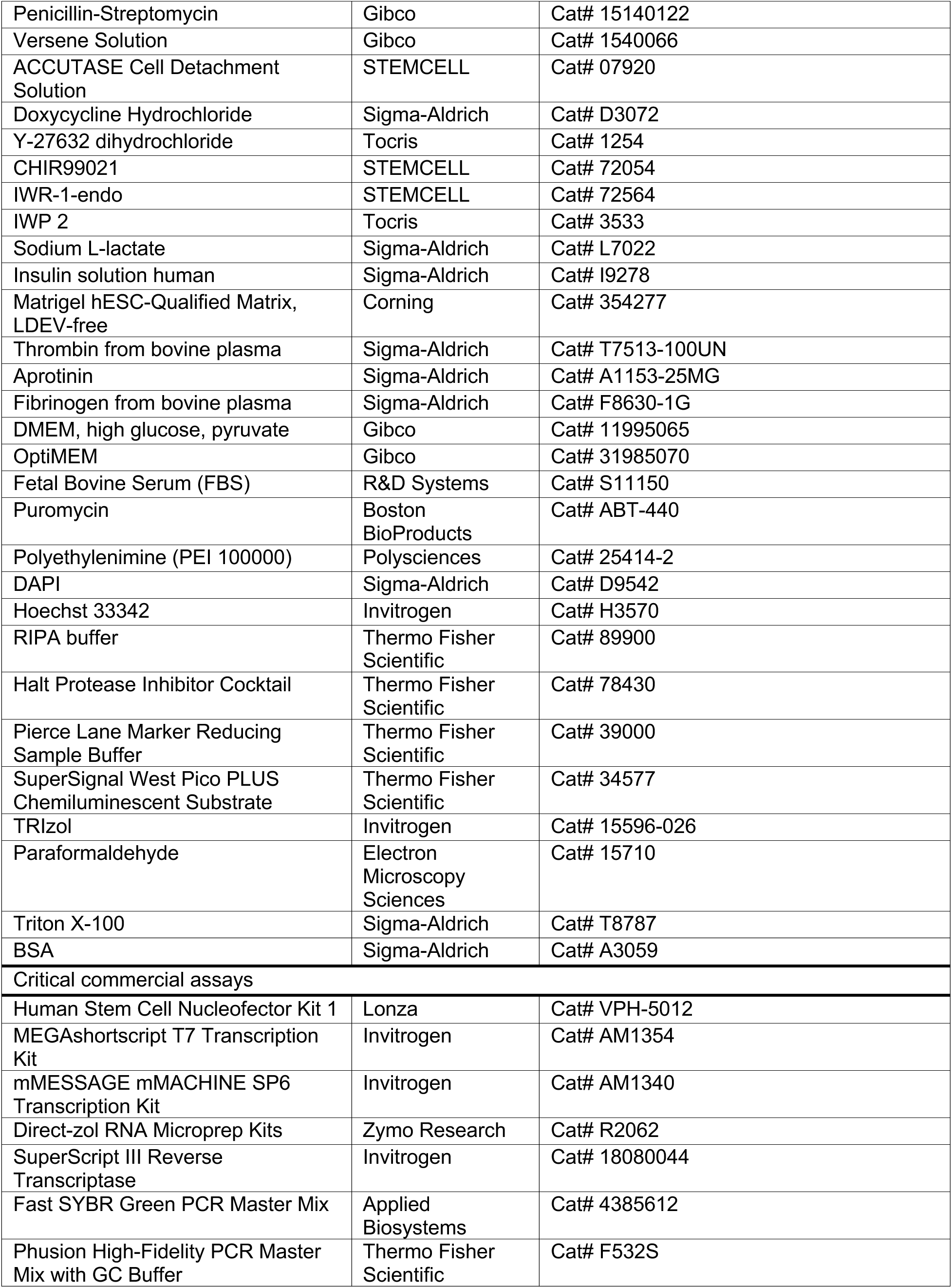
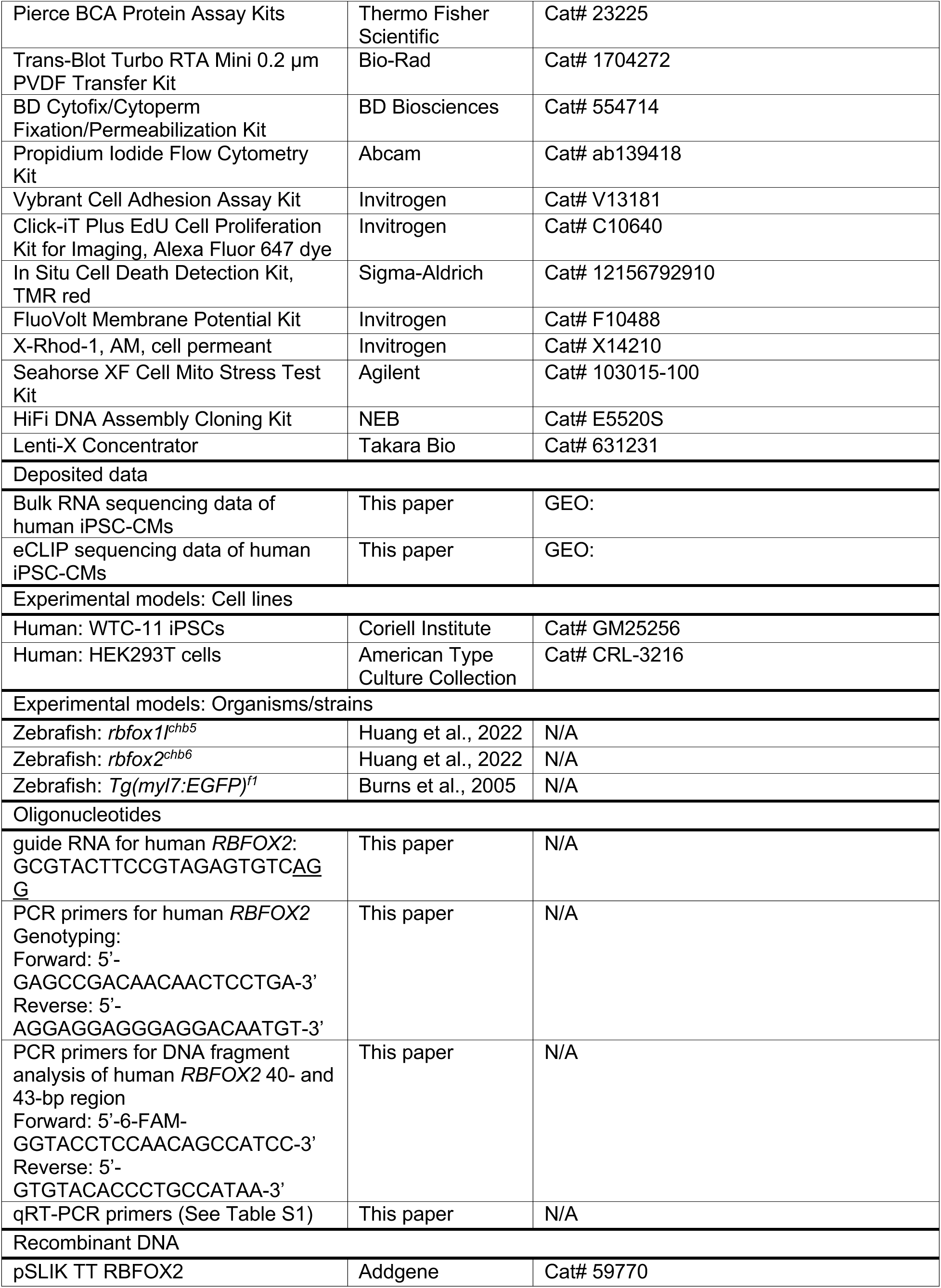
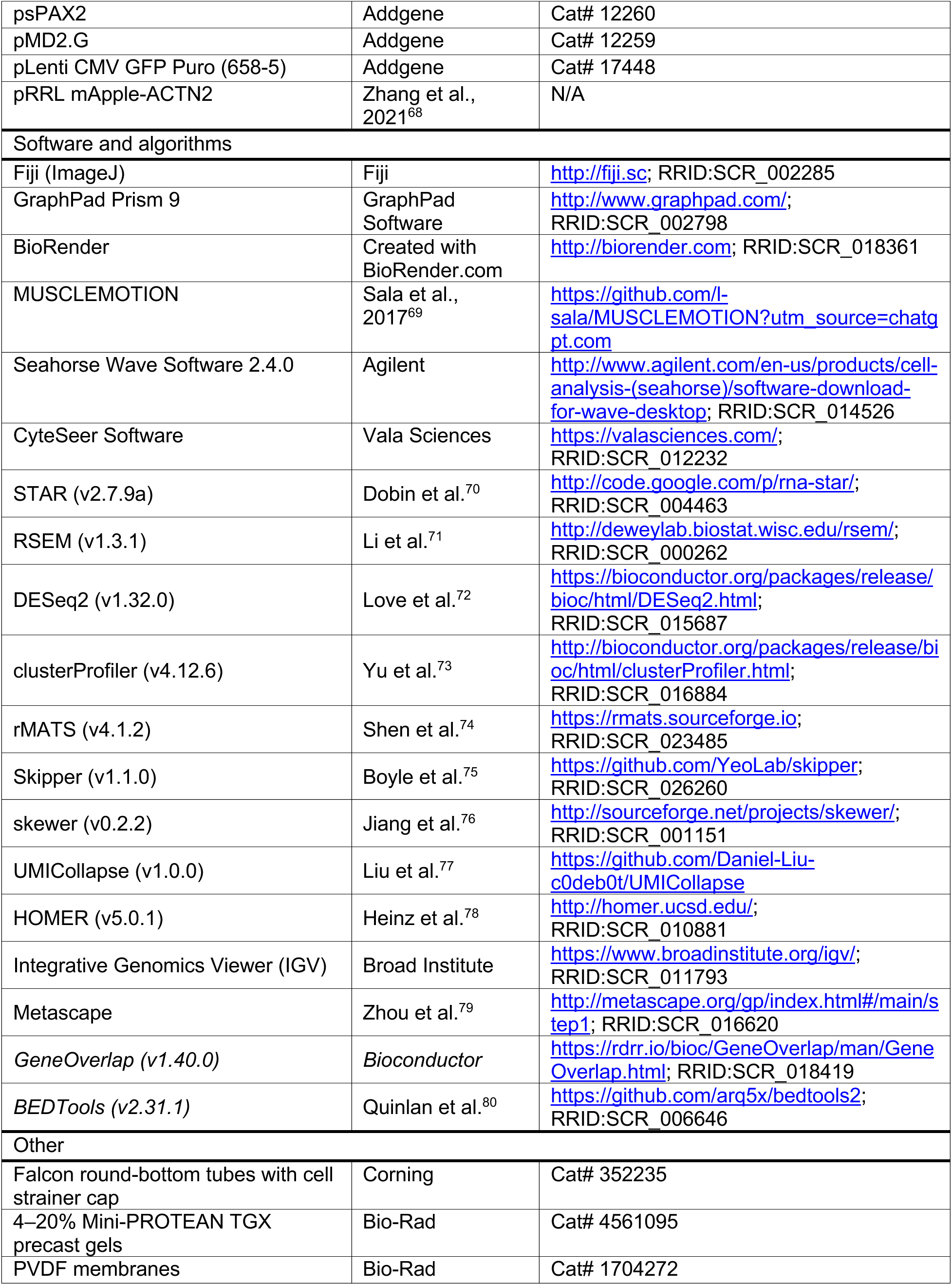
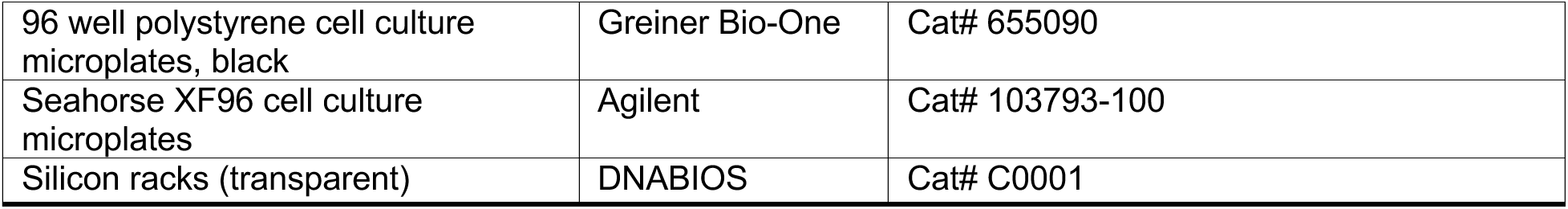

## METHODS

### Human iPSC derivation and culture

Human induced pluripotent stem cells (iPSCs) were derived from the WTC-11 background (Coriell Institute). Cells were maintained in Essential 8 Medium (Gibco) at 37°C in a humidified atmosphere with 5% CO₂ and passaged every 2-4 days using Versene (Thermo Fisher Scientific). Culture plates were pre-coated with 1% Geltrex matrix (Thermo Fisher Scientific) according to the manufacturer’s instructions.

### Zebrafish husbandry and strains

Zebrafish (Danio rerio) were maintained under standard conditions in accordance with protocols approved by the Institutional Animal Care and Use Committee (IACUC) at Boston Children’s Hospital, following the NIH Guide for the Care and Use of Laboratory Animals. The following strains were used: *rbfox1l^chb5^*, *rbfox2^chb6^*, and *Tg(myl7:EGFP)^f1^*^19,81^.

### Human iPSC genome editing

CRISPR/Cas9 genome editing was performed to generate isogenic human iPSC lines. A guide RNA (gRNA) targeting *RBFOX2* (5′-GCGTACTTCCGTAGAGTGTC-3′) was synthesized using the MEGAshortscript T7 Transcription Kit (Thermo Fisher Scientific). WTC-11 iPSCs were genetically modified to express a doxycycline-inducible SpCas9 transgene in the AAVS1 (adeno-associated virus integration site 1) safe harbor locus as previously described^82,83^. To induce Cas9 expression, WTC-11 AAVS1^Cas9/+^ hiPSCs were treated with doxycycline (2 μg/mL) for 16 h prior to transfection. Approximately 0.5 × 10⁶ cells were then nucleofected with 5 μg gRNA using the Amaxa Nucleofector system (Lonza), and plated at low density for clonal isolation. Clonal genotyping of *RBFOX2* edits was performed by PCR (Forward: 5′-GAGCCGACAACAACTCCTGA-3′; Reverse: 5′- AGGAGGAGGGAGGACAATGT-3′), followed by Sanger and amplicon sequencing to identify heterozygous and null clones (n=4 independent clones per genotype).

### Directed differentiation of hiPSCs into cardiomyocytes

Human iPSCs were differentiated into cardiomyocytes (iPSC-CMs) using a monolayer-based protocol with temporal Wnt signaling modulation, as previously described^24,84^. Cells were seeded onto 12-well plates and cultured for 48 h or until ∼50% confluence. On day 0, cells were treated with RPMI (Thermo Fisher Scientific) supplemented with B27 minus insulin (Thermo Fisher Scientific) and 6.5–7 µM CHIR99021 (STEMCELL Technologies) for 48 h. On day 2, the medium was replaced with fresh RPMI/B27(–insulin) containing 5 µM IWR-1-endo (STEMCELL Technologies) and 5 µM IWP-2 (Tocris Bioscience) for 48 h. From days 4–10, cells were maintained in RPMI/B27(–insulin) with medium changes every 48 h. On day 10, metabolic selection was performed in glucose-free RPMI supplemented with 5 mM sodium lactate (Sigma-Aldrich) for 48 h. Cells were then recovered for at least 4 days in RPMI/B27(–insulin) prior to downstream assays. For dissociation, iPSC-CMs were incubated with Accutase (Thermo Fisher Scientific) at 37 °C for 10–15 min, resuspended in RPMI/B27 supplemented with 10 µM ROCK inhibitor (Y-27632; Tocris Bioscience).

### Flow cytometry analysis

Pluripotency of hiPSCs and differentiation efficiency of hiPSC-CMs were assessed by flow cytometry. Briefly, hiPSCs and hiPSC-CMs were dissociated with Versene (5 min, 37 °C) or Accutase (10 min, 37 °C), respectively, and passed through a 70 µm cell strainer. Cells were washed and fixed using the Cytofix/Cytoperm Fixation/Permeabilization Kit (BD Biosciences) according to the manufacturer’s instructions. Fixed cells were incubated overnight at 4 °C with FITC-conjugated antibodies against SSEA-4 (Miltenyi Biotec, 130-098-371), cardiac troponin T (cTnT; Miltenyi Biotec, 130-119-575), or an IgG isotype control (Miltenyi Biotec, 130-104-611), each diluted 1:100 in wash buffer. After two washes, cells were resuspended in wash buffer and transferred to Falcon tubes equipped with cell-strainer caps (Corning). Samples were kept on ice and analyzed on a BD LSRFortessa flow cytometer (BD Biosciences).

### Quantitative real-time PCR (qRT-PCR)

Total RNA was extracted using TRIzol reagent (Invitrogen) and purified with the Direct-zol Microprep Kit (Zymo Research). cDNA was synthesized with the SuperScript III First-Strand Synthesis System (Invitrogen) following the manufacturer’s instructions. qRT-PCR was performed in triplicate on a QuantStudio 3 (Applied Biosystems) or CFX384 (Bio-Rad) instrument using the Fast SYBR Green Master Mix (Thermo Fisher Scientific). Relative mRNA expression levels were calculated using the 2⁻ΔΔCt method with GAPDH as an internal control. Isoform-specific primers were designed based on rMATS analysis to quantify alternative splicing events, whereas primers spanning constitutive exons were used to measure total transcript abundance. Primer sequences are listed in Supplementary Table S1.

### Protein immunoblotting

Samples were lysed in RIPA buffer (Thermo Fisher Scientific) supplemented with Halt Protease Inhibitor Cocktail (Thermo Fisher Scientific). Protein concentrations were determined using the Pierce BCA Protein Assay Kit (Thermo Fisher Scientific), normalized, and reduced and denatured in sample buffer (Thermo Fisher Scientific). Equal amounts of protein were separated on 4–20% Mini-PROTEAN TGX precast gels (Bio-Rad), transferred to PVDF membranes (Bio-Rad) using the Trans-Blot Turbo Transfer System (Bio-Rad), and blocked in TBS-T (Tris-buffered saline with 0.1% Tween-20, pH 7.4) containing 5% BSA. Membranes were incubated overnight at 4 °C with primary antibodies, washed with TBS-T, and incubated with HRP-conjugated secondary antibodies (Cell Signaling, 7076 and 7074) for 1 h at room temperature. Signals were detected using SuperSignal West ECL substrate (Thermo Fisher Scientific) and imaged with a GE ImageQuant LAS-4000. Images were processed and quantified using Fiji (ImageJ). Primary antibodies used were: anti-RBFOX2 (Bethyl Laboratories, A300-864A), anti–α-actinin (Abcam, ab9465), and anti-GAPDH (Cell Signaling Technology, 2118T)

### Analysis of contractility in 2D-cultured hiPSC-CMs

Contraction of 2D-cultured hiPSC-CMs in the original plates was recorded in bright-field mode (40x magnification) using a Keyence BZ-X700 microscope. Videos were captured at 33 frames per second to ensure sufficient temporal resolution for accurate quantification of contractile dynamics. For quantitative analysis of spontaneous beat frequency, the acquired time-lapse images were processed using MUSCLEMOTION, as previously described^69^.

### Cell adhesion assay

Cell adhesion was quantified using the Vybrant Cell Adhesion Assay Kit (Thermo Fisher Scientific) according to the manufacturer’s instructions. Briefly, flat-bottom 96-well plates (Greiner Bio-One) were pre-coated with 0.5% Geltrex matrix at 37 °C for 2 h. hiPSC-CMs were dissociated with Accutase (Thermo Fisher Scientific) for 10 min at 37 °C and labeled with 5 µM calcein AM (included in the kit) for 30 min at 37 °C. After two washes, cells were resuspended at 5 × 10⁵ cells/mL, and 100 µL per well was seeded into the pre-coated plates and incubated for 30 min at 37 °C. Total fluorescence was measured using a FlexStation 3 Multi-Mode Microplate Reader (Molecular Devices). After gentle washing to remove nonadherent cells, fluorescence of adherent cells was recorded. Background wells contained medium only. The percentage of adhered cells was calculated as: Cell Adhesion (%) = [(Adherent fluorescence − Background) / Total fluorescence] × 100

### Immunofluorescence, EdU staining, TUNEL staining and imaging

hiPSC-CMs were replated onto 1% Geltrex-coated 96-well plates (Greiner Bio-One) and allowed to recover for 4 days before fixation. Cells were fixed with 4% paraformaldehyde (Electron Microscopy Sciences) for 10 min, permeabilized in PBS containing 0.1% Triton X-100 for 5 min, and blocked in PBS containing 3% BSA. Cells were then incubated with primary antibodies overnight at 4 °C, followed by Alexa Fluor–conjugated secondary antibodies for 1 h at room temperature. Nuclei were counterstained with DAPI (1:1000; Sigma-Aldrich) for 5 min before imaging. EdU staining was performed using the Click-iT EdU Imaging Kit (Invitrogen) after a 2 h EdU pulse prior to fixation. TUNEL staining was performed using the In Situ Cell Death Detection Kit, TMR red (Sigma-Aldrich), followed by the same immunofluorescence procedures described above. Primary antibodies included Alpha-Actinin (ACTN2, 1:500; Abcam, ab9465) and cardiac Troponin T (cTnT, 1:500; DSHB, CT3). Secondary antibodies included goat anti-mouse IgG1 Alexa Fluor 488 (1:500; Invitrogen, A-21121), and goat anti-mouse IgG2a Alexa Fluor 555 (1:500; Invitrogen, A-21137). Images were acquired using an Olympus FV3000R resonant-scanning confocal microscope and processed with Fiji (ImageJ).

### Morphological analysis of 2D-cultured hiPSC-CMs

For cell size analysis, hiPSC-CMs were seeded into 96-well plates for 24 h and imaged under bright field (100x magnification) using the EVOS M5000 Imaging System (Thermo Fisher Scientific). Cell area was quantified from confocal images using Fiji by manually tracing individual cell boundaries with the polygon selection tool. For myofibrillar disarray analysis, immunofluorescence staining for ACTN2 followed by confocal imaging (600x magnification) was performed as above. Myofibrillar disarray was quantified in Fiji by assessing the alignment consistency of sarcomeric Z-discs across multiple regions of interest, as previously described^85^. All analyses were performed in a blinded manner.

### Transmission electron microscopy (TEM)

hiPSC-CMs were prepared and processed, and ultrathin sections were imaged at the Harvard Medical School Electron Microscopy Facility as previously described^19^. Briefly, cells were seeded onto Aclar coverslips in 24-well plates and allowed to recover for 4 days, then fixed for 1 h at room temperature in a 1:1 mixture of RPMI/B27(–insulin) and FGP fixative (2.5% paraformaldehyde, 5% glutaraldehyde, and 0.06% picric acid in 0.2 M cacodylate buffer). Samples were post-fixed in 1% osmium tetroxide/1.5% potassium ferrocyanide, stained with 1% uranyl acetate in maleate buffer (pH 5.2), dehydrated through a graded ethanol series, transitioned through propylene oxide prior to resin infiltration, and embedded in Taab 812 resin (Marivac-Canemo). Ultrathin sections (80 nm) were cut using a Leica Ultracut S ultramicrotome, post-stained with 0.2% lead citrate, and imaged using either a Philips Tecnai BioTwin Spirit or a JEOL 1200× transmission electron microscope.

### Seahorse analyzer analysis of oxygen consumption rate (OCR)

Mitochondrial function in hiPSC-CMs was assessed using the Seahorse XF Cell Mito Stress Test Kit (Agilent) according to the manufacturer’s instructions. hiPSC-CMs were seeded at 7.5 × 10⁴ cells per well onto 1% Geltrex-coated Seahorse XF96 cell culture microplates (Agilent). Cells were maintained in RPMI/B27(–insulin) medium for 4 days to allow recovery prior to the assay. Oxygen consumption rate (OCR) was recorded using a Seahorse XF96 Extracellular Flux Analyzer, and data were analyzed with Wave software (version 2.4.0).

### Calcium transient and membrane potential analysis

Calcium and membrane voltage imaging was recorded on a Vala Sciences Kinetic Image Cytometer (KIC) as previously described with some modifications^86^. Briefly, 1 × 10⁵ hiPSC-CMs were seeded onto black-walled 96-well plates (Greiner Bio-One) and cultured for 4 days. Cells were incubated for 30 min at 37°C with a dye-loading solution prepared in Tyrode’s buffer (137 mM NaCl, 2.7 mM KCl, 1 mM MgCl₂, 1.8 mM CaCl₂, 0.2 mM Na₂HPO₄, 12 mM NaHCO₃, 5.5 mM D-glucose, pH 7.4) containing X-Rhod-1 (1:2000, Thermo Fisher Scientific), FluoVolt membrane potential dye (1:1000) with PowerLoad (1:100, Thermo Fisher Scientific), and Hoechst 33342 (10 μg/mL, Invitrogen). After two washes with Tyrode’s buffer, cells were incubated in fresh buffer for 10 min prior to imaging. Temperature-controlled recordings were acquired over 30 s using a three-phase protocol: 10 s of spontaneous activity, 10 s of pacing at 1 Hz and 15 V (via MyoPacer stimulator, IonOptix), and 10 s of post-pacing recovery. Line-scan images were processed using a custom ImageJ macro, and quantification of calcium transient kinetics and membrane potential dynamics was performed using CyteSeer software (Vala Sciences).

### Generation and functional assessment of engineered heart tissues (EHTs)

EHTs were generated as previously described^44^, with minor modifications. Briefly, 1 × 10⁶ hiPSC-CMs were used per construct. Three days after casting, cells were transduced with an adenovirus encoding channelrhodopsin-2 fused to GFP (Ad-ChR2-GFP, a gift from Vassilios J Bezzerides)^87^. EHTs were maintained in RPMI/B27(–insulin), supplemented with 0.1% insulin (Sigma-Aldrich), 5 µg/mL aprotinin (Sigma-Aldrich), and 1% penicillin–streptomycin (Gibco), with medium changes every other day. EHTs were cultured in 24-well plates and kept in a stage-top incubator at 37°C with 5% CO₂. For functional assessment, spontaneous and paced contractions were recorded starting on day 7, and quantitative analyses were performed on day 28. Optical pacing was applied using blue LEDs positioned above the wells, and contractions were recorded from below at 30 frames per second through a 561-nm long-pass filter (Semrock BLP02-561R-32) using an 8-mm f/1.4 lens (Thorlabs MVL8M1) mounted on a Basler acA1920 camera. Post deflection was tracked using the Multi-Template Matching plugin in Fiji (ImageJ), and twitch force was calculated by beam-bending analysis based on the known Young’s modulus of the posts, as described elsewhere^88^.

### DNA fragment analysis

DNA fragment analysis was performed using cDNA synthesized as described above and Phusion DNA Polymerase (Thermo Fisher Scientific) with 6-FAM–labeled primers (listed in the Oligonucleotides table). PCR products were purified and analyzed at the MGH CCIB DNA Core Facility, where fluorescently labeled DNA fragments were separated on an ABI 3730xl Genetic Analyzer (Applied Biosystems). Fragment sizes and relative abundances were determined using GeneMapper v6.0 software (Applied Biosystems).

### Plasmid cloning

For overexpression experiments, the mApple–ACTN2 fusion construct was PCR-amplified from the pRRL-mApple–α-actinin-2 plasmid (a gift from Christopher S. Chen)^68^. The early isoform of *RBFOX2* containing the 40-bp C-terminal exon was amplified from the pSLIK-TT-RBFOX2 plasmid (Addgene). To generate the late *RBFOX2* isoform (containing the 43-bp exon), the truncated isoform (containing both the 40-bp and 43-bp exons), and a 3×FLAG control construct, oligonucleotide pairs encoding the respective sequences were synthesized by Integrated DNA Technologies (IDT). All constructs were PCR-amplified and cloned into the pLenti-CMV backbone (Addgene) using the NEBuilder HiFi DNA Assembly Kit (New England Biolabs). Plasmids were propagated in NEB competent *E. coli* according to the manufacturer’s instructions.

### Lentivirus production

Lentivirus was produced by transfecting HEK293T cells (ATCC CRL-3216) seeded in 150-mm dishes with 20 mL DMEM (Thermo Fisher Scientific) supplemented with 10% FBS (R&D Systems). Cells were transfected with 28 μg of lentiviral transfer vector, 21 μg psPAX2 (Addgene), and 7 μg pMD2.G (Addgene) using polyethyleneimine (PEI; Polysciences) in Opti-MEM (Thermo Fisher Scientific) at a PEI:DNA ratio of 3:1 (w:w). Medium was replaced 24 h post-transfection, and virus-containing supernatant was harvested at 72 h. Lentivirus was concentrated using Lenti-X Concentrator (Takara) according to the manufacturer’s instructions. To determine viral titers, hiPSCs were transduced with serial dilutions of the concentrated virus and selected with 1 μg/mL puromycin (Boston BioProducts). Transduction efficiency was quantified by counting puromycin-resistant colony-forming units.

### Lentivirus overexpression studies

Lentiviral overexpression constructs were transduced into hiPSCs at a multiplicity of infection (MOI) >1. Medium was replaced the following day. For rescue overexpression experiments, transduced hiPSCs were selected for single-cell clones, and successful integration of the transgene was confirmed by Sanger sequencing. Verified clones were subsequently expanded and differentiated into hiPSC-CMs as described above for downstream functional analyses.

### Zebrafish genotyping and mRNA microinjection

Germline transmission of CRISPR/Cas9-induced *rbfox1l* and *rbfox2* deletions was detected using fluorescent PCR and DNA fragment analysis as previously described^19^. Microinjection of mRNA into zebrafish embryos was performed as described previously^19^. Human *RBFOX2* early (containing the 40-bp C-terminal exon), late (containing the 43-bp C-terminal exon), and truncated (containing both the 40-bp and 43-bp C-terminal exons) isoforms, were generated from constructs described in the plasmid cloning section. Each isoform was subcloned into the pCS2+ vector. Plasmids were linearized with *SacII* and used as templates for in vitro transcription using the mMESSAGE mMACHINE SP6 Transcription Kit (Thermo Fisher Scientific). Approximately 1 nL of mRNA (∼100 pg) was microinjected into zebrafish embryos at the one-cell stage.

### Bulk RNA-seq library preparation and sequencing

Total RNA was isolated and purified as described above. Biological replicates included three control, five *RBFOX2* heterozygous, and five null samples, each comprising 1–2 × 10⁶ hiPSC-CMs from independent differentiations. Poly(A)-selected RNA-seq libraries were prepared using a standard Illumina mRNA library preparation workflow. Sequencing was performed on an Illumina NovaSeq 6000 platform (2 × 150 bp paired-end, single index), yielding approximately 350 million paired-end reads per lane.

### Differential gene expression analysis

Raw FASTQ files were first processed for quality control and aligned to the reference genome (GENCODE v38) using STAR (v2.7.9a)^70^. Gene-level read counts in STAR outputs were obtained for each sample and imported into DESeq2 (v1.32.0) for differential expression analysis^72^. Significance was defined by an adjusted p-value (False Discovery Rate, FDR) < 0.05 and an absolute fold change > 1.5. Transcripts per million (TPM) were also calculated using RSEM (v1.3.1) to provide normalized expression measures for visualization and cross-sample comparison^71^. Gene Ontology (GO) enrichment analysis was performed using clusterProfiler (v4.12.6)^73^.

### Alternative splicing analysis

Alternative splicing event (ASE) analysis of bulk RNA-seq data was performed using rMATS tools (v4.1.2)^74^, applying an FDR < 0.05 to identify the significant events. rMATS detects five categories of ASEs, including skipped exon (SE), alternative 5′ splice site (A5SS), alternative 3′ splice site (A3SS), mutually exclusive exons (MXE), and retained intron (RI). Functional enrichment analysis of ASE-associated genes was performed as described above for GO term enrichment.

### Integration analysis of alternative splicing across datasets

For integrative analysis of ASE between *RBFOX2* het and control samples and during human iPSC-CM differentiation (day 7 vs. day 14), bulk RNA-seq data from a published study were reanalyzed^21^. The dataset included hiPSC-CM samples collected at day 7 and day 14 (n = 3 biological replicates per time point). The same ASE analysis pipeline described above was applied to identify differential splicing events. Overlapping ASEs across datasets were defined based on shared genomic coordinates of regulated exons and inclusion level changes, and functional enrichment of the overlapping genes was analyzed for enriched GO terms as described above.

### eCLIP-seq library preparation

Large-scale cultures (∼1.5 × 10⁷ cells) of wild-type hiPSC-CMs (day 15) were harvested from an Eppendorf DASbox bioreactor as previously described^89^. eCLIP was performed following established protocols^90,91^. Cells were resuspended in minimal ice-cold PBS and UV-crosslinked on ice (254 nm, 400 mJ cm⁻²) to covalently link RNA–protein complexes. Crosslinked cells were scraped, pelleted (200 × g, 3 min), washed with PBS, flash-frozen on dry ice, and stored at −80 °C. Immunoprecipitations were carried out with anti-RBFOX2 antibody (Bethyl Laboratories, A300-864A) or IgG control (Thermo Fisher Scientific, 02-6102). RNA fragments bound to RBFOX2 were ligated to adapters containing random barcodes (N5/N10), reverse-transcribed, and PCR-amplified to generate sequencing libraries. Libraries were sequenced on an Illumina NovaSeq 6000 platform (single-end 50 bp). Two biological replicates were sequenced per condition.

### eCLIP-seq analysis

Sequencing reads were processed using the Skipper pipeline^75^. Adapters were trimmed with skewer^76^, reads were aligned to the human genome (hg38) using STAR^70^, and PCR duplicates were removed with UMICollapse^92^. Crosslinking sites were identified by counting 5′ read ends within fixed-size tiled windows across annotated genic regions. Windows were stratified by GC content to estimate and correct for GC bias. Enrichment of RBFOX2 binding was determined by comparing immunoprecipitated reads with their size-matched input (SMinput) controls. Motif analysis was performed on significant peaks (*P* < 0.001 and >8-fold change) using HOMER^78^ , as previously described^91^. Genome browser tracks were visualized in IGV^93^, and pathway enrichment analysis was performed using Metascape^79^.

### Overlap analysis between RBFOX2 binding and alternative splicing events

RBFOX2 eCLIP-seq peaks were determined from processed data and converted to BED format. Genes containing RBFOX2-bound regions were compared with differentially spliced genes using *GeneOverlap* (https://bioconductor.org/packages/GeneOverlap/). Overlap between RBFOX2 binding peaks and alternatively spliced exons or splice junctions (±250 nt) was assessed using *BEDTools*^80^. Fisher’s exact test was used to evaluate enrichment, and overlaps with an odds ratio > 1.5 and *P* < 0.001 were considered significant.

## QUANTIFICATION AND STATISTICAL ANALYSIS

Statistical analyses were performed using GraphPad Prism 9. The number of biological replicates, statistical tests, and exact *P* values are provided in the corresponding figure legends. Data are presented as mean ± standard deviation (SD) or as median with interquartile range, as indicated. Comparisons between two groups were performed using unpaired two-tailed *t* tests. Comparisons among three or more groups were analyzed by one-way ANOVA followed by Tukey’s multiple comparisons test, and when two independent variables were assessed, two-way ANOVA with Bonferroni’s post hoc test was applied. Sample sizes were not predetermined using statistical methods. Randomization and investigator blinding were not applied unless otherwise specified due to the nature of the study.

