## Supplementary material for "Dosage-sensitive *RBFOX2* autoregulation promotes cardiomyocyte differentiation by maturing the transcriptome": Manuscript For bioRxiv_v2

### SUPPLEMENTAL INFORMATION

**Figures S1, Generation and characterization of *RBFOX2*-deficient human iPSC-CMs; related to Figure 1**

**Figure S2, *RBFOX2* deficiency causes altered structural and electrophysiological properties in human iPSC-CMs; related to Figure 2**

**Figure S3, *RBFOX2* regulates alternative splicing of transcripts encoding sarcomere, cytoskeletal and adhesion components in human iPSC-CMs; related to Figure 4**

**Figure S4, Overexpression of *RBFOX2* isoforms and ACTN2 in human iPSC-CMs; related to Figures 6 and 7.**

**Figure S5, Isoform-specific and ACTN2 rescue of *RBFOX2* haploinsufficiency restores contractile kinetics in engineered heart tissues; related to Figures 6 and 7.**

**Table S1 Sequence of primers used in this study.**

**Video S1. 2D-cultured iPSC-CMs, related to Figure 1**
Video microscopy recording of 2D-cultured iPSC-CMs at day 15 from *RBFOX2* CTRL, het and null groups. Scale bar, 500 µm.

**Video S2. 3D-cultured EHTs, related to Figure 4**
Video microscopy recording of 3D-cultured EHTs at day 28 under 1hz pacing from *RBFOX2* CTRL, het and null groups. Scale bar, 1 mm.

**Spreadsheet S1, DEGs and GO terms, related to Figure 3**
(1) Gene expression list in het vs CTRL
(2) Gene expression list in null vs CTRL
(3) GO (Biological Process) terms enriched among down-regulated DEGs in het vs. CTRL
(4) GO (Biological Process) terms enriched among down-regulated DEGs in null vs. CTRL
(5) GO (Biological Process) terms enriched among up-regulated DEGs in het vs. CTRL
(6) GO (Biological Process) terms enriched among up-regulated DEGs in null vs. CTRL

**Spreadsheet S2, ASEs and GO terms, related to Figure 3**
(1) Differential alternative splicing events (A3SS) in het vs. CTRL
(2) Differential alternative splicing events (A5SS) in het vs. CTRL
(3) Differential alternative splicing events (MXE) in het vs. CTRL
(4) Differential ASEs (RI) in het vs. CTRL
(5) Differential ASEs (SE) in het vs. CTRL
(6) GO (Biological Process) terms enriched among Differential ASEs in het vs CTRL
(7) Differential ASEs (A3SS) in null vs. CTRL
(8) Differential ASEs (A5SS) in null vs. CTRL
(9) Differential ASEs (MXE) in null vs. CTRL
(10) Differential ASEs (RI) in null vs. CTRL

(11) Differential ASEs (SE) in null vs. CTRL
(12) GO (Biological Process) terms enriched among Differential ASEs in null vs CTRL

**Spreadsheet S3, eCLIP targets and GO terms, related to Figure 3**
(1) eCLIP enriched peaks
(2) GO (Biological Process) terms enriched in eCLIP target genes

**Spreadsheet S4, eCLIP+ASE+ genes and GO terms, related to Figure 4**
(1) eCLIP+ASE+ genes in het vs. CTRL
(2) GO (Biological Process) terms enriched in eCLIP+ASE+ genes in het vs. CTRL
(3) eCLIP+ASE+ genes in null vs. CTRL
(4) GO (Biological Process) terms enriched in eCLIP+ASE+ genes in null vs. CTRL

**Spreadsheet S5: Overlapping ASEs and GO terms**, **related to Figure 4**
(1) Overlapping ASEs between Het vs. CTRL and Day7 vs. Day14 from Lau et al., Cell Rep. 2019
(2) GO (Biological Process) terms enriched in genes with significant overlapping ASEs between het vs. CTRL and Day7 vs. Day14

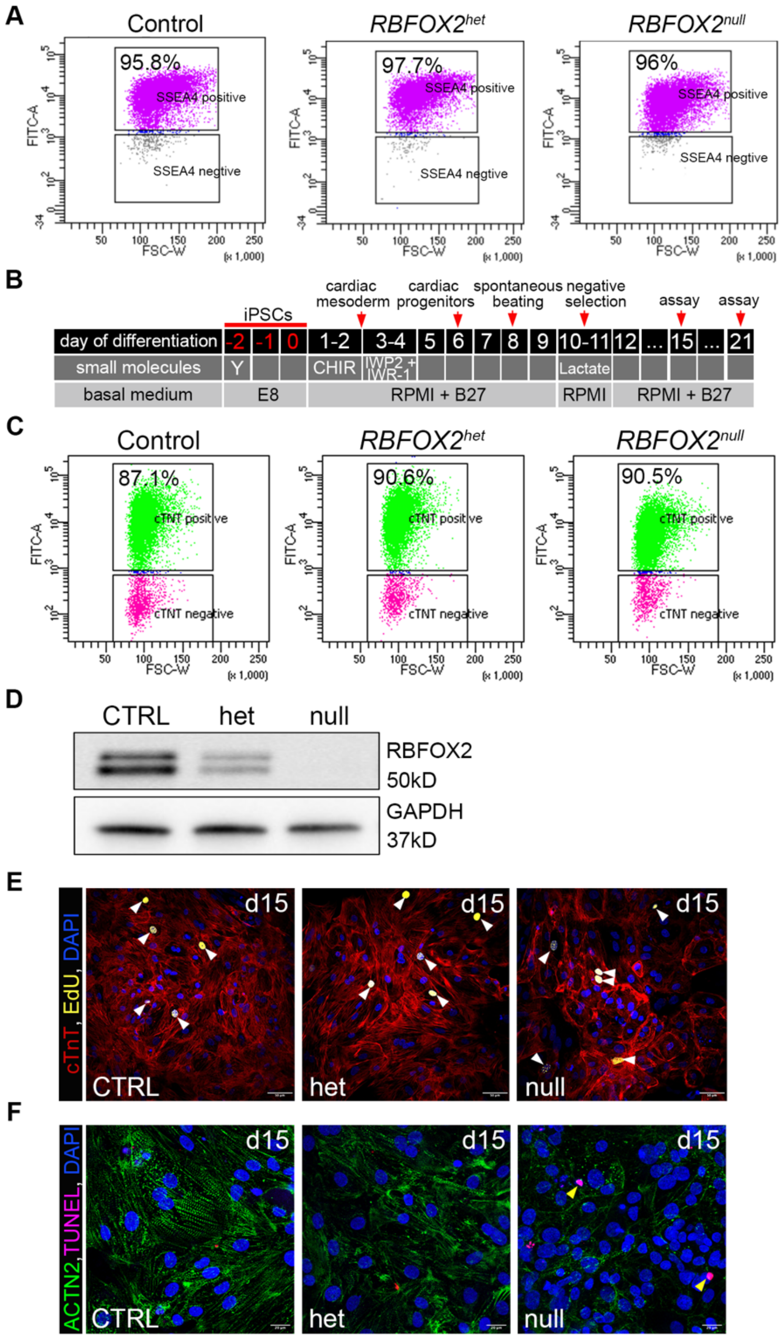


**Figure S1. Generation and characterization of *RBFOX2*-deficient human iPSC-CMs.**

(A) Flow cytometry analysis of pluripotency marker SSEA4 in CTRL, *RBFOX2* het and *RBFOX2* null iPSCs (n = 3 biological replicates).

(B) Schematic of the directed differentiation protocol from iPSCs to cardiomyocytes, indicating timing of small molecule treatments, basal media, and negative selection.

(C) Flow cytometry analysis of cardiac marker cTnT in CTRL, het and null iPSC-CMs at day 21. Numbers indicate the percentage of cTnT-positive cells shown in green; cTnT-negative cells are shown in magenta (n = 3).

(D) Western blot analysis of RBFOX2 protein levels in CTRL, het, and null iPSC-CMs at day 21, with GAPDH as loading control (n = 3).

(E) Immunofluorescence staining of CTRL, het, and null iPSC-CMs at day 15 for cTnT (red), EdU (yellow), and DAPI (blue) (n = 3). Arrowheads indicate EdU-positive CMs. Scale bar, 50 μm.

(F) Immunofluorescence staining of CTRL, het, and null iPSC-CMs at day 15 for ACTN2 (green), TUNEL (magenta), and DAPI (blue) (n = 3). Arrowheads indicate TUNEL-positive cardiomyocytes. Scale bar, 20 μm.

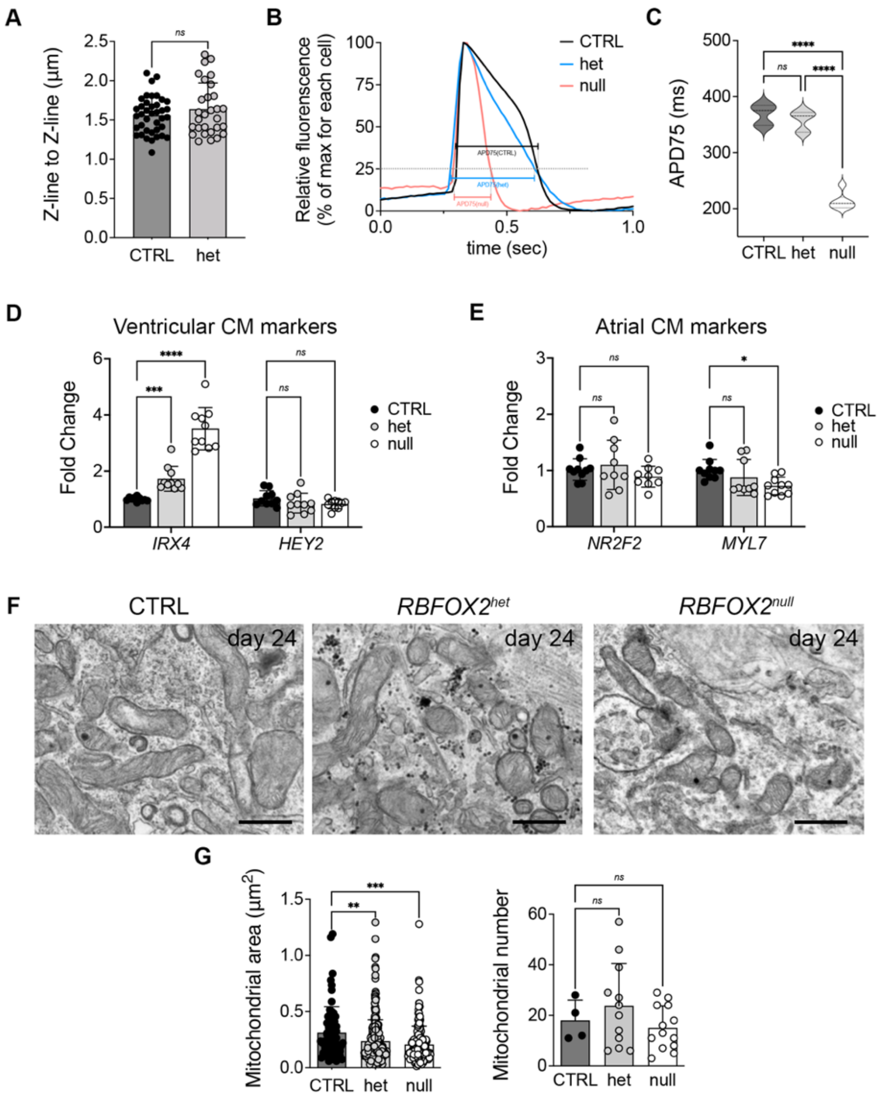


**Figure S2.** ***RBFOX2*** **deficiency causes altered structural and electrophysiological properties in human iPSC-CMs.**

(A) Quantification of Z-line to Z-line distance in day 24 CTRL, het, and null iPSC-CMs from transmission electron micrographs (n = 3; each dot represents one sarcomere).

(B, C) Representative action potential traces under 1 Hz pacing (B) and quantification of action potential duration at 75% repolarization (APD75; C) in day 21 CTRL, het, and null iPSC-CMs (n = 3).

(D, E) qRT-PCR analysis of ventricular cardiomyocyte markers (*IRX4, HEY2*; D) and atrial cardiomyocyte markers (*NR2F2, MYL7*; E) in day 21 CTRL, het, and null iPSC-CMs (n = 9-10).

(F, G) Transmission electron microscopy images of mitochondria (F) and quantification of mitochondrial area (each dot represents one mitochondrion) and mitochondrial number (each dot represents one image count) (G) in day 24 CTRL, het, and null iPSC-CMs (n = 3).

Data are presented as mean ± SD (A, C-E, and G). n indicates biological replicates. Statistical analyses were performed using unpaired two-tailed t-test (A) or one-way ANOVA with Tukey's post hoc test (C-E, G). Significance levels: *$p < 0.05$; **$p < 0.001$; ***$p < 0.001$; ****$p <$ 0.0001; *ns*, no [illegible]

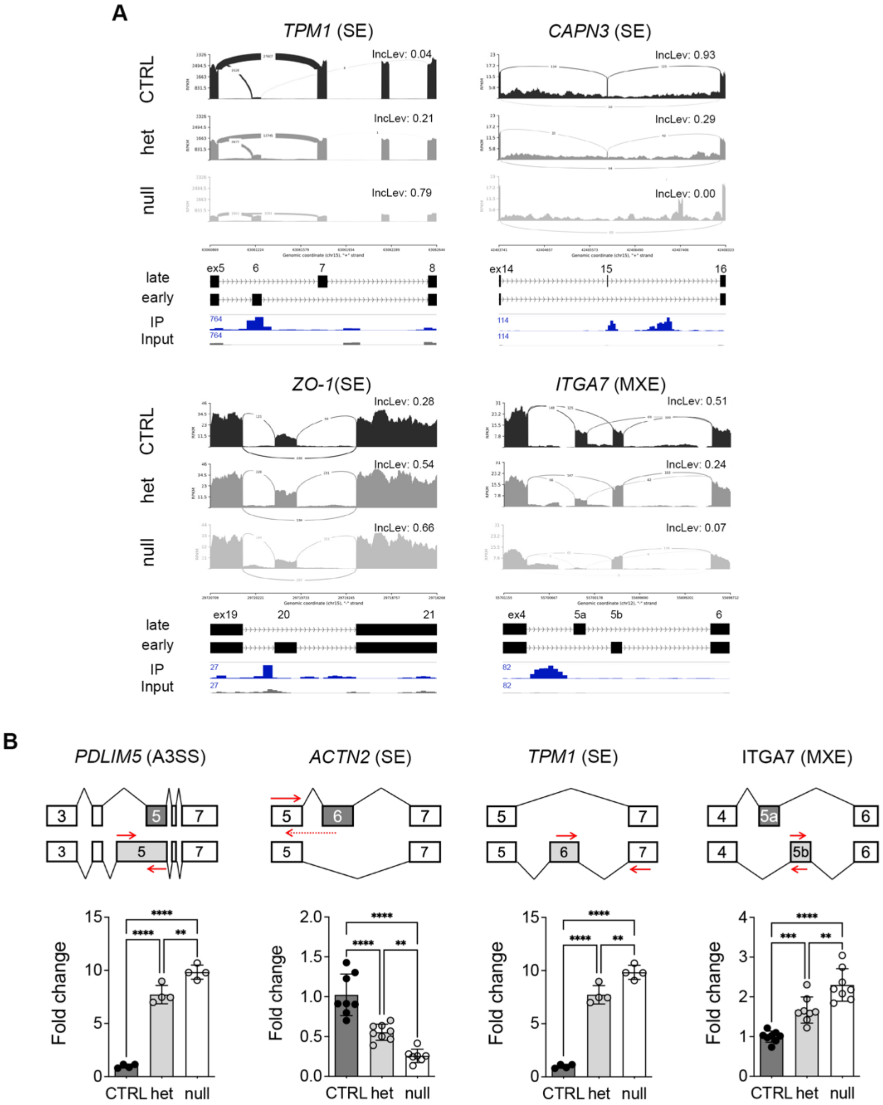


**Figure S3.** ***RBFOX2*** **regulates alternative splicing of transcripts encoding sarcomere, cytoskeletal and adhesion components in human iPSC-CMs.**

(A) Sashimi plots of alternative splicing events in selected RBFOX2 targets (*TPM1, CAPN3, ZO-1,* and *ITGA7*) from Bulk RNA seq of day 21 CTRL and het iPSC-CMs, with representative RBFOX2 eCLIP binding peaks shown. Inclusion level differences (IncLev) are indicated. Numbers on tracks indicate normalized read counts. SE, skipped exon; MXE, mutually exclusive exon; ex, exon; early, early isoform; late, late isoform.

(B) qRT-PCR validation of selected alternative splicing events in CTRL, het, and null iPSC-CMs (n = 4-8), including *PDLIM5* (A3SS), *ACTN2* (SE), *TPM1* (SE), and *ITGA7* (MXE). Splicing schematics are shown above each quantification plot. Red arrows indicate primer positions used to amplify specific exons. Isoform expression levels were normalized to total transcript levels. A3SS, alternative 3′ splice site; SE, skipped exon; MXE, mutually exclusive exon.

Data are presented as mean ± SD (B). n indicates biological replicates. Statistical analyses were performed using one-way ANOVA with Tukey's post hoc test. Significance levels: $**p < 0.01$; $***p < 0.001$; $****p < 0.0001$; *ns*, not significant.

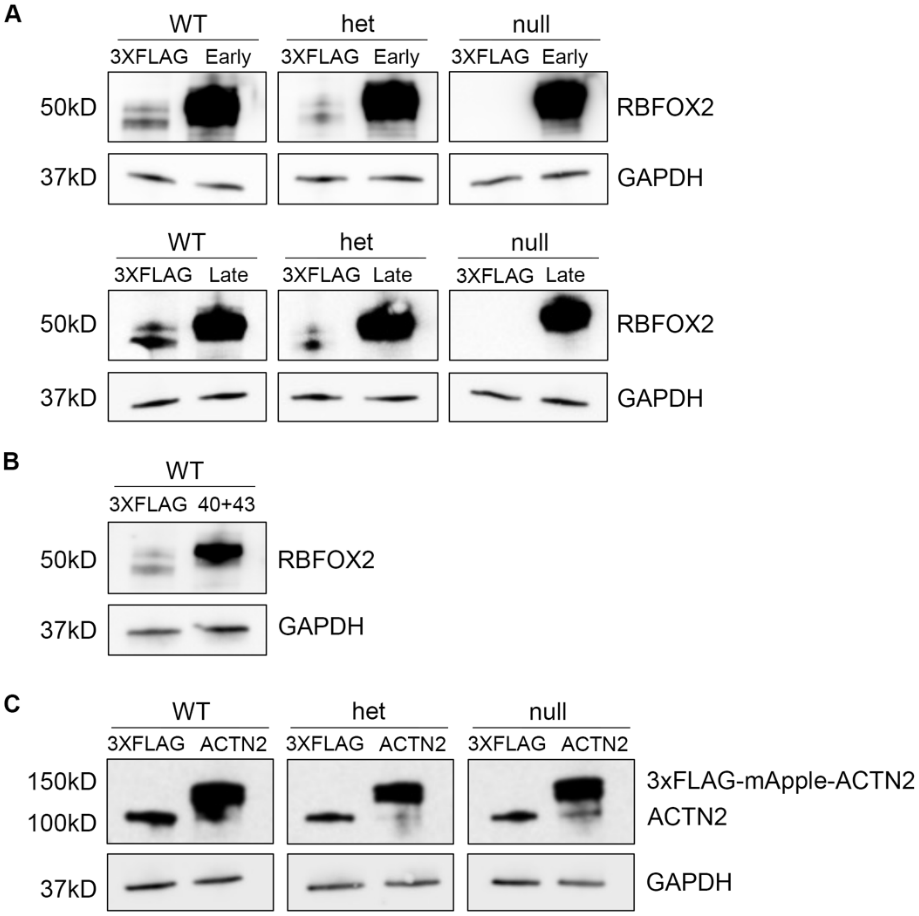


**Figure S4. Overexpression of *RBFOX2* isoforms and ACTN2 in human iPSC-CMs.**

(A-C) Western blot analysis of lentiviral overexpression in day 21 iPSC-CMs. RBFOX2 expression was assessed in WT, het, and null iPSC-CMs transduced with lentivirus encoding 3×FLAG control, Early, or Late *RBFOX2* isoforms (A), and in WT iPSC-CMs transduced with lentivirus encoding 3×FLAG control or the aberrant 40+43 *RBFOX2* isoform (B). ACTN2 expression was analyzed in WT, het, and null iPSC-CMs transduced with lentivirus encoding 3×FLAG control or 3×FLAG-mApple-ACTN2, showing detection of both exogenous 3×FLAG-mApple-ACTN2 and endogenous ACTN2 (C). GAPDH served as a loading control in all experiments. n = 3 biological replicates.

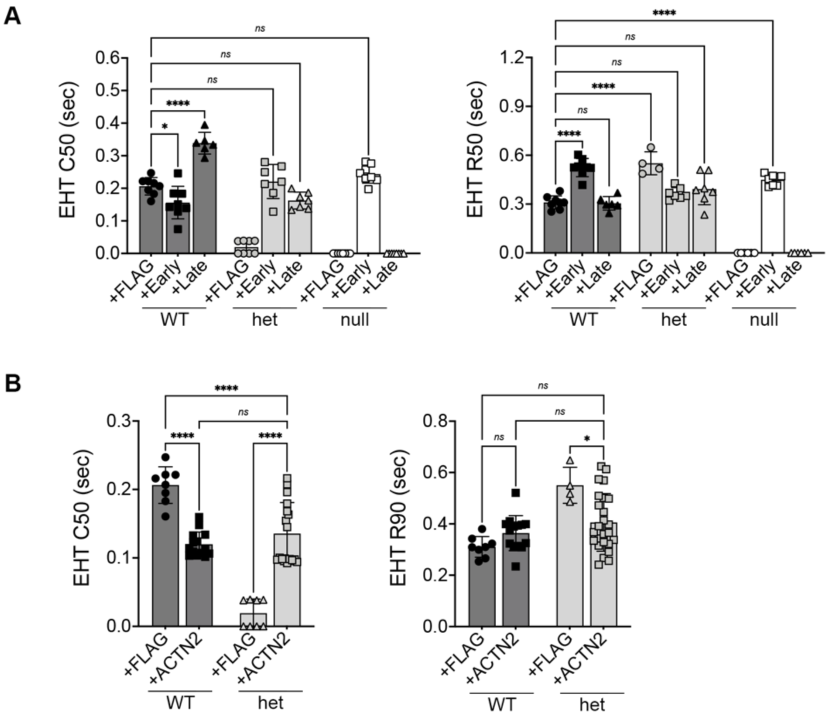


**Figure S5. Isoform-specific and ACTN2 rescue of *RBFOX2* haploinsufficiency restores contractile kinetics in engineered heart tissues.**

(A, B) Quantification of contraction time to 50% peak (C50) and relaxation time to 50% peak (R50) in engineered heart tissues (EHTs under 1 Hz pacing, day 28) generated from WT, het, and null iPSC-CMs transduced with lentivirus encoding 3×FLAG control, Early, or Late *RBFOX2* isoforms (A) and with FLAG or ACTN2 (B) (n = 3; each dot represents one EHT).

Data are presented as mean ± SD (A and B). n indicates biological replicates. Statistical analyses were performed using one-way ANOVA with Tukey's post hoc test. Significance levels: *$p$ < 0.05; ****$p$ < 0.0001; *ns*, not significant.

**Table S1 Sequence of primers used in this study.**

| **Primers for qPCR** | | |
|---|---|---|
| Gene Name/Target Region | Forward Primer | Reverse Primer |
| GAPDH | TGAGTACGTCGTGGAGTCCA | AGAGGGGGCAGAGATGATGA |
| RBFOX2 | CAGATGTTTGGGCAGTTTGG | CTCGATTTTACGGCCCTCTA |
| MYH6 | CTTCAACCACCACATGTTCG | GGCTTCTGGAAATTGTTGGA |
| MYH7 | GACGGAGGAGGACAGGAAAA | TCCTCATTCAAGCCCTTCGT |
| IRX4 | TTGGACTCCTGGGAACATGGACAA | ATGCTTCAGGGTATCTGGCCTCTT |
| HEY2 | GAAAGAGCCGCTAGGAGCAG | GCTAGTACTTTGCCCCGAGT |
| NR2F2 | GCAAGTGGAGAAGCTCAAGG | GCTTTCCACATGGGCTACAT |
| MYL7 | CCGTCTTCCTCACGCTCTT | TGAACTCATCCTTGTTCACCAC |
| PDLIM5 | TGTAGAGGAGAAAGGAGCCC | GCTTTCCACAGGCTACACAC |
| PDLIM5 (longer exon 5 region) | ATCACATGCTTCCCCTTCAC | CCCTCTGCTAGCTCCTGAGA |
| ACTN2 | GGCACCCAGATTGAGAACAT | CCCCTTTGCTGGCTATGTAA |
| ACTN2 (exon 5 and 6 region) | GGCAACGTGAAAATGACCCT | AGGCCATCTTTCCAGCTAGT |
| TPM1 | AGCCGACGTAGCTTCTCTGA | TCATCTTTTTGGGCTCGACT |
| TPM1 (exon 6 and 7 region) | CCAAGTCCGACAGCTGGA | CTGAGCCTCCAGTGACTTCA |
| ITGA7 | ACCAATACCCTGACCTGCTG | CTATAGCTGCTGGGGACTGC |
| ITGA7 (exon 5b region) | GGTTGCTTTTTGTGACCAAC | GCTATTGAGGGCCAAGTCTC |
